# MALT1 protease inhibition restrains glioblastoma progression by reversing tumor-associated macrophage-dependent immunosuppression

**DOI:** 10.1101/2024.09.26.614808

**Authors:** Juliana Hofstätter Azambuja, Saigopalakrishna S. Yerneni, Lisa M. Maurer, Hannah E. Crentsil, Gabriela N. Debom, Linda Klei, Mei Smyers, Chaim T. Sneiderman, Kristina E. Schwab, Rajesh Acharya, Yijen Lin Wu, Prasanna Ekambaram, Dong Hu, Pete J. Gough, John Bertin, Ari Melnick, Gary Kohanbash, Riyue Bao, Peter C. Lucas, Linda M. McAllister-Lucas

## Abstract

MALT1 protease is an intracellular signaling molecule that promotes tumor progression via cancer cell-intrinsic and cancer cell-extrinsic mechanisms. MALT1 has been mostly studied in lymphocytes, and little is known about its role in tumor-associated macrophages. Here, we show that MALT1 plays a key role in glioblastoma (GBM)-associated macrophages. Mechanistically, GBM tumor cells induce a MALT1-NF-κB signaling axis within macrophages, leading to macrophage migration and polarization toward an immunosuppressive phenotype. Inactivation of MALT1 protease promotes transcriptional reprogramming that reduces migration and restores a macrophage “M1-like” phenotype. Preclinical *in vivo* analysis shows that MALT1 inhibitor treatment results in increased immuno-reactivity of GBM-associated macrophages and reduced GBM tumor growth. Further, the addition of MALT1 inhibitor to temozolomide reduces immunosuppression in the tumor microenvironment, which may enhance the efficacy of this standard-of-care chemotherapeutic. Together, our findings suggest that MALT1 protease inhibition represents a promising macrophage-targeted immunotherapeutic strategy for the treatment of GBM.

**Graphical abstract.:** The effects of tumor cell-induced CARD9-BCL10-MALT1 (CBM) activation (left) and MALT1 protease inhibition (right) on GBM associated macrophages in the tumor microenvironment.
Cartoon of cellular components of a GBM tumor with an immunosuppressive TME characterized by “M2-like macrophages” (left) and conversion to a more immune-reactive tumor microenvironment characterized by “M1-like macrophages and increased effector T-cells (right) as a result of MALT1 protease inhibition.

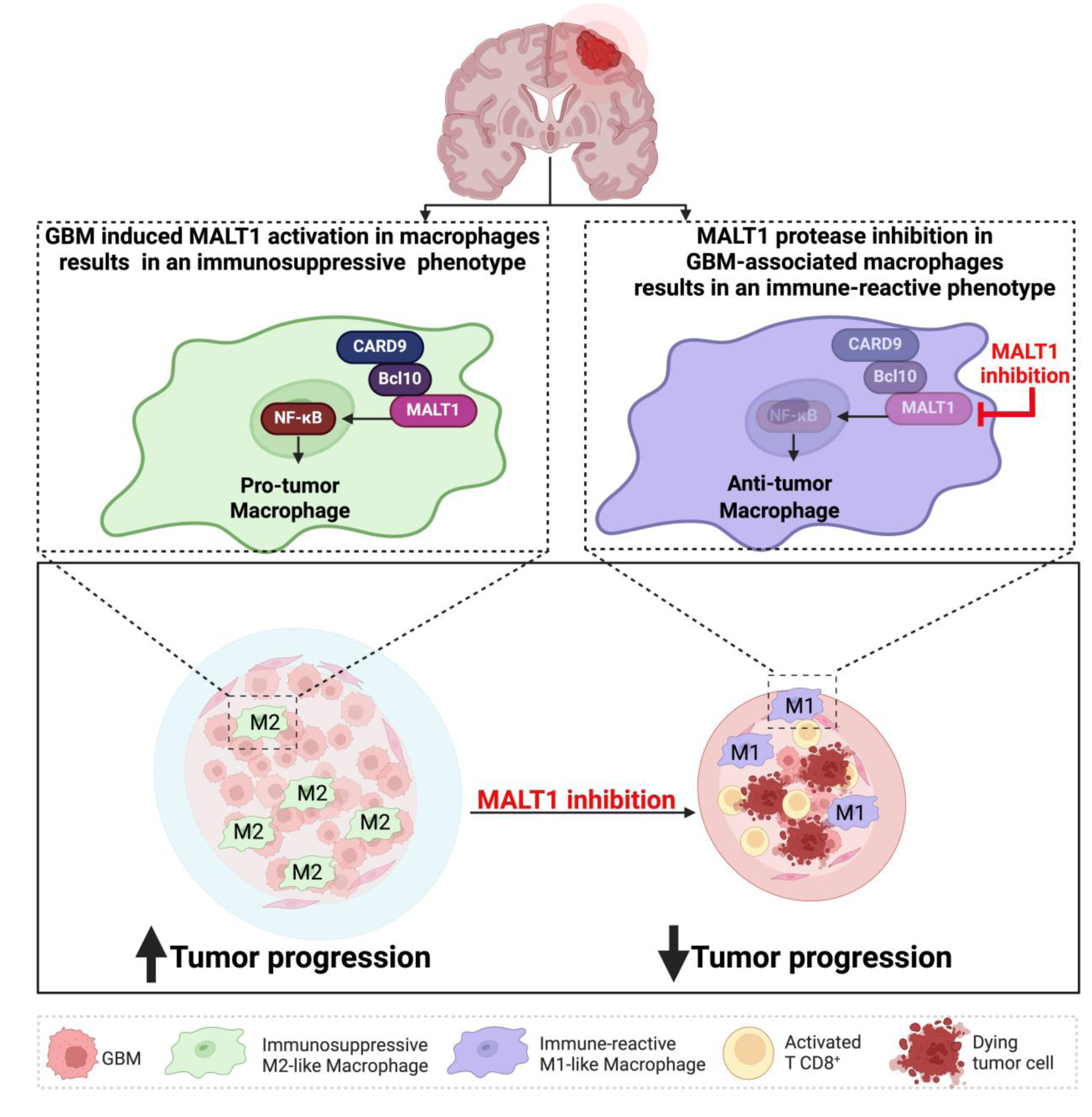

## Introduction

Glioblastoma (GBM) is the most common and aggressive primary malignant brain tumor in adults, with a median overall survival of about 14 months^1^. GBM remains incurable and inevitably recurs after conventional cytotoxic treatments including radiation and chemotherapy^2,3^. GBM is one of the most challenging cancers to treat, largely owing to tumor heterogeneity, the delicate location of the tumors limiting surgical options, the impermeability of the blood-brain barrier (BBB) to many therapeutic agents, the pro-tumorigenic influence of the tumor microenvironment (TME), and the long-term complications of treatment, including significant neurologic sequelae that limit treatment intensity^4^. There is an urgent unmet need for improved strategies to treat GBM.

The TME plays a critical role in promoting GBM invasion and progression by stimulating angiogenesis and suppressing anti-tumor immune responses^5^. In GBM, tumor-associated bone-marrow derived macrophages and brain-resident microglia, together referred to as “TAMs”, represent the major immune cell populations within the TME, accounting for up to half of the cells of the tumor mass^6^. GBM TAMs usually demonstrate an immunosuppressive (“M2-like”) phenotype that promotes cancer growth^7^. TAMs therefore represent a potential target for GBM immunotherapy. Although it is well established that macrophages play essential roles in driving immune suppression and supporting tumor progression, the molecular mechanisms that regulate the tumor-promoting functions of macrophages remain only partially understood. Thus, elucidating the molecular mechanisms that regulate GBM TAM polarization has the potential to inform new therapeutic avenues.

Here, we demonstrate the critical role of MALT1 in promoting GBM pathogenesis via its role in macrophages in the TME. MALT1 is the downstream effector protein of the CARMA-CARD/BCL10/MALT1 (CBM) intracellular signaling complex, and it functions via at least two mechanisms: (1) MALT1 acts as a scaffold to bind and recruit downstream signaling proteins and (2) MALT1 acts as a protease to enzymatically cleave and inactivate multiple specific substrate proteins^8^. The CBM complex plays a key role in normal adaptive immunity by mediating antigen receptor-dependent activation of the pro-survival/pro-inflammatory NF-κB transcription factor in T and B cells, leading to lymphocyte cytokine production and proliferation^9^. Deregulated CBM activation is considered a hallmark of multiple subtypes of lymphoma, where gain-of-function mutations in CBM components or upstream regulators of the CBM complex drive constitutive MALT1-dependent NF-κB activity, resulting in tumor progression^10^. Thus, MALT1 is considered a promising new therapeutic target in lymphoma and MALT1 inhibitors are currently being evaluated in Phase I clinical trials for the treatment of lymphoid malignancies^11^.

Beyond lymphoma, MALT1 may also represent a promising therapeutic target in other cancers due to the potential dual benefit of inhibiting MALT1 proteolytic activity within both tumor cells and lymphocytes. First, inappropriate/deregulated activation of CBM signaling, resulting in constitutive MALT1 protease activity, is now recognized as a common feature in an increasing number of solid tumors including certain lung cancers, breast cancers and others, and inhibiting CBM signaling within these cancer cells abrogates their proliferation and survival^12–17^. Second, selective inhibition of MALT1 protease activity, without inhibiting MALT1 scaffolding activity, has been shown to induce an inflammatory response^18–21^. This effect of MALT1 protease inhibition has been thus far attributed to a relative loss of Treg activity and/or reprogramming of immunosuppressive Treg cells into IFN-γ-secreting effector cells. These alterations in Treg and T effector activity caused by MALT1 protease inhibition have been shown to result in enhanced T effector cell anti-tumor immunity^22–24^. Due to the multi-faceted role of MALT1 in both tumor cells and in lymphocytes, there is considerable interest in leveraging MALT1 protease inhibition as an innovative approach to cancer therapy.

Thus far, investigation of MALT1’s influence on tumor immune response has focused on its role in T cells. Here, we report that MALT1 proteolytic activity plays a key role in macrophages by regulating tumor associated macrophage polarization in GBM. Specifically, we find that MALT1 promotes an immunosuppressive M2-like macrophage phenotype in GBM. Furthermore, we demonstrate that MALT1 protease activity promotes macrophage infiltration into the GBM tumor mass. Using a combination of genetic and pharmacologic approaches, we show that MALT1 protease inhibition abrogates tumor associated macrophage-mediated immune suppression and promotes an anti-tumor immune response in GBM.

## Results

### Selective and specific MALT1 protease inhibition has minimal impact on GBM cancer cell phenotypes

Several recent reports have suggested a potential role for MALT1 in promoting GBM tumor progression^12,14,25,26^. Specifically, these reports showed that inhibiting MALT1, either by genetic knockout or pharmaceutical inhibition, reduces GBM cancer cell survival, proliferation, migration, and invasion. These studies combined *in vitro* analyses of cancer cell behavior with *in vivo* immunodeficient GBM xenograft mouse models. In light of these findings, we sought to further investigate the mechanisms by which MALT1 influences GBM pathogenesis. To begin, we demonstrated that members of the CBM complex, CARMA3, BCL10 and MALT1, are present in both mouse and human GBM cancer cells (Fig. S1a,b). Further, GBM cells of both species demonstrate constitutive MALT1 proteolytic activity, as indicated by cleavage of the MALT1 substrate, N4BP1, and this can be effectively abrogated by siRNA-mediated MALT1 knockdown (Fig. S1a,b).

Interestingly, we found that while MALT1 knockdown had a substantial negative effect on clonogenic potential, particularly for human GBM cells, there was only a modest negative effect on either mouse or human cell viability and no effect whatsoever on cell migration (Fig. S1a,b). We then evaluated the impact of specifically blocking MALT1 protease activity with MLT-748, a relatively new and highly selective small molecule MALT1 protease inhibitor^27^. Results were generally similar to what we observed with MALT1 knockdown, except that clonogenic activity of human GBM cells was not impacted (Fig. S1c,d).

Because the effects of MLT-748 on GBM cell phenotype were generally modest, we compared MLT-748 to first-generation MALT1 inhibitors (MI-2 and mepazine) which had been used in the earlier publications on GBM and found that only MI-2 elicits a major effect on GBM cancer cell viability (Fig. S2). It is possible that this difference relates to the fact that MI-2 is a covalent irreversible inhibitor while mepazine and MLT-748 are both allosteric, reversible MALT1 protease inhibitors^28^. This unique effect of MI-2, in comparison to newer and more specific MALT1 protease inhibitors such as MLT-748, may also be related to what is now understood to be its significant off-target profile^29–31^. Taken together, our findings suggest that while MALT1 does contribute to some cancer cell intrinsic properties of GBM, the specific inhibition of MALT1 protease activity alone may not have a major impact on the neoplastic cells themselves.

### In GBM, MALT1 is most highly expressed in tumor-associated myeloid cells

With recent reports indicating that MALT1 protease inhibition can elicit anti-tumor lymphocyte re-activation in cancer^31^, we next decided to investigate the role of MALT1 in the GBM TME. To begin, we analyzed data from The Cancer Genome Atlas (TCGA) to query the expression of *MALT1* mRNA in specific GBM subtypes. Our initial analyses revealed that levels of *MALT1* mRNA are significantly elevated in Grade IV GBM tumors when compared with non-tumor brain tissue samples (Fig. 1a). We found that among GBM tumor subtypes, *MALT1* expression is higher in mesenchymal GBM, a molecular subtype that represents approximately 30- 50% of GBM cases^32^, in comparison to the classical or proneuronal subtypes (Fig. 1b). Notably, mesenchymal GBM is considered the most aggressive subtype, tends to have the worst overall survival rates compared to other subtypes and is often characterized by an extensive inflammatory infiltrate^32^. We also found that higher *MALT1* mRNA expression in GBM tumors is associated with lower probability of survival (Fig. 1c). Of note, a survival advantage for patients with “*MALT1*-low expression” GBM tumors was similarly observed by Jacobs *et al* using a different dataset, thus corroborating our findings^12^.

**Fig. 1.**
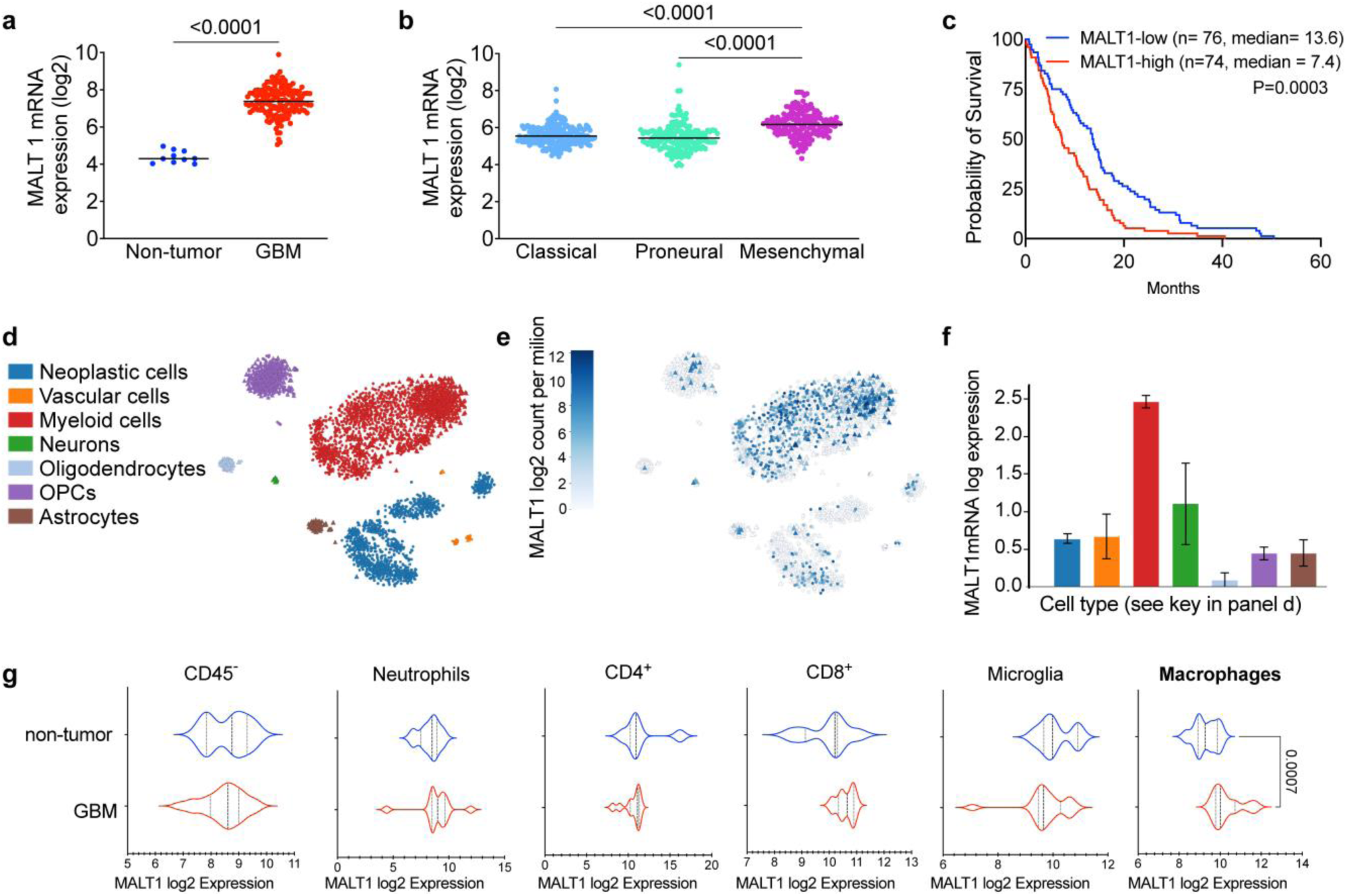
MALT1 is expressed in TAMs in GBM. GlioVis data portal was used to analyze brain tumor mRNA expression datasets. **(a)** Comparison of *MALT1* mRNA expression in GBM tumor samples with non-tumor brain samples. Each dot represents one clinical sample. **(b)** Comparison of *MALT1* mRNA expression in 3 different GBM molecular subtypes **(c)** Probability of survival of 150 patients diagnosed with GBM, grouped by high (top 50%) or low (bottom 50%) *MALT1* mRNA level. **(d)** 2D-tSNE representation of single cells in the GBM TME (graph generated in http://gbmseq.org). **(e)** *MALT1* mRNA expression across different cell types isolated from GBM tumors (n=6 individuals, GSE84465 dataset) **(f)** Bar graph showing *MALT1* expression in the different cell subsets found in the GBM samples. Cell types are indicated by colors as shown on the left in panel D. **(g)** Representation of MALT1 expression across different cell types within GBM tumor core or non-tumor tissue, evaluated by single-cell RNA-sequencing. Data in b were analyzed by 1-way ANOVA, followed by Tukey’s multiple-comparisons. Statistical significance in a and g was determined by two-tailed Student’s t-test. Significant p values are indicated in the figure.

To evaluate MALT1 expression in specific cells within the GBM TME, we analyzed *MALT1* mRNA using single-cell RNA-sequencing (RNA-seq) data from a cohort of 6 individual patients diagnosed with GBM^33^. Marker-based cell-type annotation revealed 7 distinct clusters comprising the major cell types present in the GBM TME, including malignant (neoplastic) cells, vascular cells, myeloid cells (macrophages, microglia, dendritic cells and neutrophils), neurons, oligodendrocytes, oligodendrocyte precursor cells (OPCs) and astrocytes (Fig. 1d). Our analysis showed that, as previously reported^34^, the myeloid cluster represents the most prevalent cell type in the GBM TME. Among the TME cells, *MALT1* was most abundantly expressed in myeloid cells (Fig. 1e,f). Single-cell RNA-seq comparison between normal brain and GBM tumor tissue in a separate dataset^35^ showed that *MALT1* expression is higher in tumor-associated macrophages compared to healthy, non-tumor associated macrophages (Fig. 1g, right). While MALT1 activity within T cells has been shown to influence the tumor immune response to certain solid tumors, namely colon cancer and melanoma,^23,24^, the role of MALT1 in GBM TAMs has not yet been investigated. The prevalent expression of *MALT1* in the myeloid compartment suggests that MALT1 activity may affect the function of myeloid cells in the GBM TME.

### MALT1 protease inhibition prevents macrophage polarization toward an M2-like phenotype in response to GBM cells

We next sought to investigate the impact of MALT1 inhibition on GBM-associated myeloid cells. First, we confirmed that macrophages express CBM complex components CARD9, BCL10 and MALT (Fig. S3a). Of note, while members of the Caspase Recruitment Domain coiled coil (CARD-CC) protein family, CARMA1 and CARMA3, serve as the scaffold for the CBM complex in lymphoid cells and epithelial cells, respectively, CARD9 fulfills this role in myeloid cells^36^. We also demonstrated that MALT1 is proteolytically active in these cells and that pharmacologic treatment with MLT-748 abrogates this proteolytic activity (Fig. S3b). Interestingly, we noticed that MLT-748 treatment reduces CARD9 and MALT1 levels in these macrophages; the mechanism by which this occurs and the impact of this reduction on downstream signaling are not known.

Next, we tested the effect of co-culturing macrophages with GBM tumor cells. First, we examined the effect of co-culture on the MALT1-NF-κB signaling axis in macrophages. For these studies, we utilized a RAW264.7 mouse macrophage cell line that was engineered to stably express a MALT1 protease activity reporter. In these cells, cleavage of the engineered MALT1 protease substrate site results in luciferase activity^37^ allowing us to quantify MALT1 protease activity within macrophages and compare this activity in the absence or presence of co-cultured GBM cancer cells. We observed that co-culture with syngeneic GBM cells induces MALT1 protease activity within the macrophages by 3-fold (Fig. 2a). As expected, MLT-748 blocks this GBM tumor cell- induced MALT1 protease activity in the macrophages. We also evaluated the activation of NF-κB in macrophages in response to co-culture with GBM tumor cells using the RAW-Blue macrophage cell line (InvivoGen), which stably expresses an NF-κB inducible secreted embryonic alkaline phosphatase (SEAP) reporter gene. We found that co-culture with GBM cells induces a 10-fold increase in NF-κB activity in these macrophages (Fig. 2b). Pharmaceutical blockade of either CARD9, the CARD-CARMA family member that is a component of the CBM complex within macrophages ^38^, or of MALT1 abrogates this response, indicating that the CBM complex mediates GBM tumor cell-induced NF-κB activation in macrophages. For these studies, we utilized BRD5529, a protein-protein interaction inhibitor that prevents CARD9-BCL10-MALT1 complex activity by disrupting the interaction between CARD9 and the ubiquitin ligase TRIM62, thereby inhibiting CARD9-dependent signaling by preventing its ubiquitinylation^39^.

**Fig. 2.**
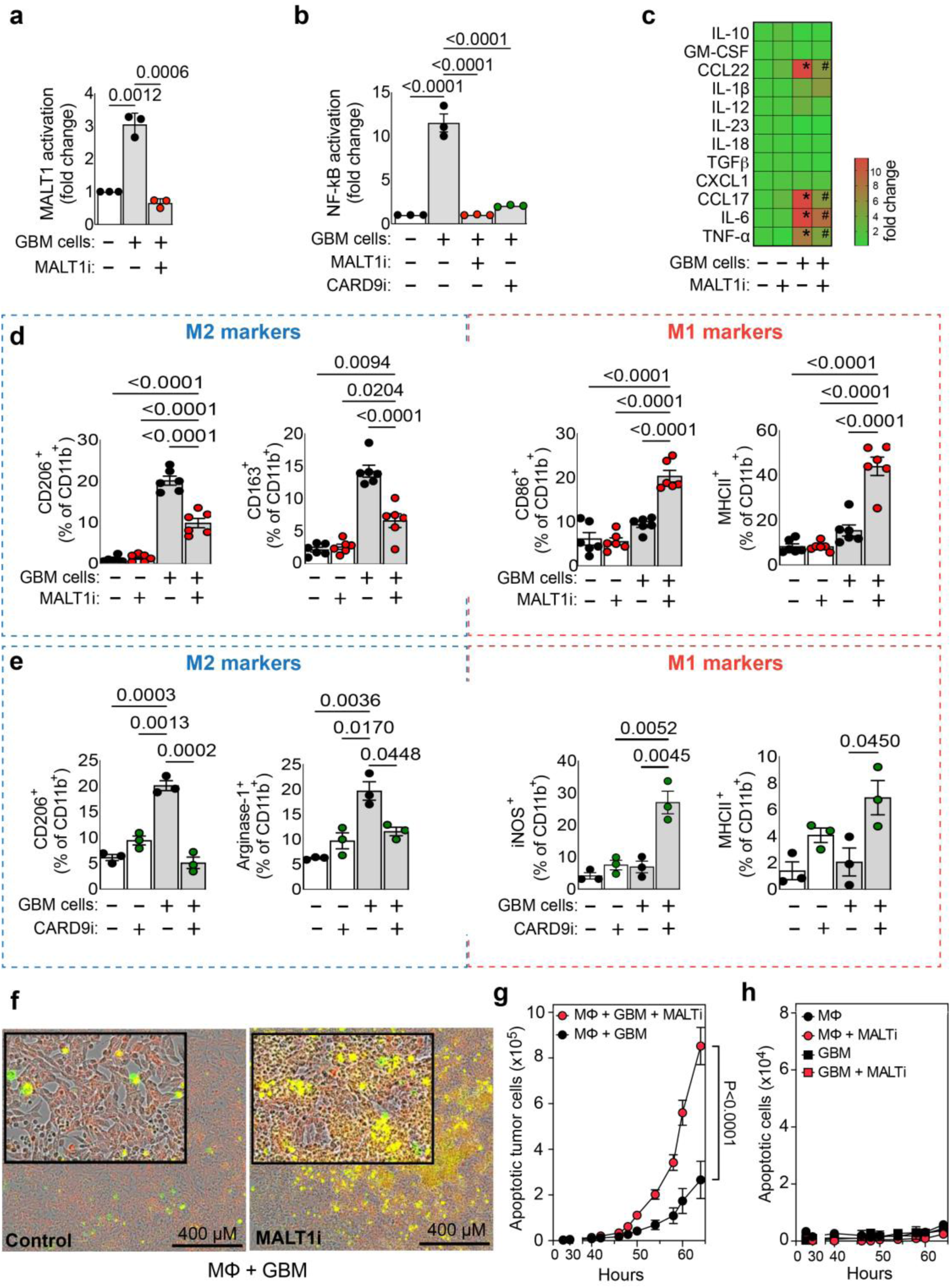
Pharmacologic inhibition of CARD9/BCL10/MALT1 complex activity reprograms M2-like GBM-associated macrophages into M1-like macrophages. **(a)** Induction of MALT1 protease activity or **(b)** NF-κB activity in RAW264.7 macrophages co-cultured with GL261-GBM cells. Macrophages were pretreated for 2 hours with or without the MALT1-protease inhibitor MLT-748 (MALT1i, 5 µM) or the CARD9 inhibitor BRD5529 (CARD9i, 50 µM) prior to co-culture. MALT1-GloSensor luciferase reporter macrophages and RAW-Blue macrophages which harbor an NF-κB-inducible secreted embryonic alkaline phosphatase reporter gene were used to measure MALT1 protease or NF-κB activity, respectively, after 48 hours of co-culture. Results are shown as fold change relative to control macrophages cultured alone. **(c)** Cytokines and chemokines in the cell culture supernatant were measured by cytometric bead array 48 h after co-culturing primary mouse macrophages with GL261-GBM cells in the absence or presence of MALT1i. Cytokine levels are presented as a heat map indicating fold change relative to the control macrophages alone. *Significant difference compared to macrophage control alone. ^#^Significant difference compared to macrophage + GBM co-culture. **(d,e)** Changes in macrophage M1/M2-like polarization markers after co-culture with GBM cells. (d) RAW264.7 mouse macrophages were co-cultured with GL261 mouse GBM cells in the absence or presence of MLT-748 (MALT1i, 5 µM). (e) RAW264.7 macrophages were cocultured with GL261 cells in the absence or presence of BRD5529 (CARD9i, 50 µM). Macrophages within the co-cultures were identified by gating for CD11b positive cells. **(f-h)** The Incucyte® live-cell analysis system was used to measure tumor cell killing in real-time. A dual color monitoring system tracked tumor cell killing by using fluorescently labeled tumor cells (red-Cell Tracker) and caspase-3/7 reagent (green) to track apoptosis. (f) Fluorescent images of RAW264.7 macrophages co-cultured with GBM-GL261 tumor cells -/+ MALT1i. Apoptotic tumor cells are indicated by yellow. (g) Co-cultures of mouse RAW264.7 macrophages with GL261-GBM cells were performed to compare macrophage-dependent tumor cell apoptosis in the absence/presence of MALT1i. (h) Controls demonstrate that MALT1i has no effect on apoptosis of GL261-GBM tumor cells or RAW264.7 macrophages when either are cultured alone. All values are represented as mean ± SD. Data were analyzed by 1-way ANOVA, followed by Tukey’s multiple-comparisons. P values are indicated in the figure.

Studies have shown that NF-κB is required to maintain the tumor-promoting phenotype of tumor-associated macrophages and that NF-κB activity in myeloid cells promotes an overall immunosuppressive TME^40^. Thus, our next step was to evaluate whether the MALT1-NF-κB axis within macrophages could be targeted to block the tumor-promoting functions of macrophages and to restore anti-tumor activity. First, we found that when co-cultured with GBM cancer cells, macrophages indeed assumed a protumor/M2-like phenotype, defined by enhanced release of CCL22, CCL17, IL-6 and TNF-α (Fig. 2c) and increased expression of both CD206 and CD163 (Fig. 2d, left) in response to their exposure to GBM cells. Of note, GBM cells cultured alone demonstrate undetectable secretion of these cytokines (not shown). Importantly, this M2 phenotypic response was abrogated by concomitant MALT1 protease inhibition with MLT-748, which resulted in a reduction of CCL22, CCL17, IL-6 and TNF-α secretion (Fig. 2c), along with a decrease in M2-like CD206^+^ and CD163^+^ expression on macrophages (Fig. 2d, left). MALT1 protease inhibition simultaneously enhanced expression of the M1 markers, CD86 and MHCII, on the GBM-associated macrophages (Fig. 2d, right), supporting the notion that MALT1 inhibition can induce GBM-associated macrophages to undergo an M2-like→M1-like phenotypic conversion. As a control, we confirmed that MLT-748 treatment has no effect on expression of CD206, CD163, CD86 and MHCII on GBM cells cultured alone (Fig. S4a).

We also evaluated the effect of CARD9 inhibitor BRD5529 on macrophage/GBM co- cultures. Analysis of RAW264.7 murine macrophages co-cultured with murine GL261 GBM cells showed that like MALT1 inhibition, CARD9 inhibition also abrogates GBM cell-induced M2-like macrophage polarization and promotes an M1-like phenotypic switch (Fig. 2e). Interestingly, we found that neither MLT-748 nor BRD5529 had a major effect on the expression of most M1/M2 markers on macrophages when these cells were cultured alone (Fig. 2d,e). Together, these results suggest that the impact of pharmacologic CBM complex inhibition on macrophage polarization appears to occur specifically in the context of GBM cell exposure. Therefore, this effect of MALT1 protease or CARD9 inhibition *in vivo* might only be detected within the GBM TME.

In addition, we tested whether these observations in mouse GBM-associated macrophages are also recapitulated in human GBM-associated macrophages. To obtain human macrophages, we isolated CD14^+^ cells from the buffy coat of three different healthy donors and then generated primary macrophages according to established protocols^41^. Flow cytometric analyses showed that when co-cultured with human GBM cells, human primary macrophages similarly undergo M2- like immunosuppressive macrophage polarization and that pretreatment of the macrophages with MLT-748 abrogates this effect of the GBM tumor cells (Fig. S4b, left), similar to what is seen in the mouse macrophage/GBM cell co-culture experiments. Also similar to mouse GBM/macrophage co-cultures, MLT-748 pretreatment induces an increase in markers associated with M1-like immune-activated anti-tumor macrophage phenotype in these human co-cultures (Fig. S4b right).

### MALT1 protease inhibition endows macrophages with an enhanced capacity to kill GBM tumor cells

We next sought to determine the effect of MALT1 protease inhibition on macrophage-dependent killing of cancer cells. For these studies, we developed a tumor cell killing assay using fluorescently labeled cancer cells (red) labeled with Caspase-3/7 Dye (green) to detect cancer cells undergoing apoptosis (Fig. 2f). This assay measures yellow fluorescence intensity (green/red mix) over time to quantify actively dying cancer cells, using the automated Incucyte real-time imaging system. Using this assay, we found that MLT-748 treatment enhances RAW264.7 macrophage-dependent killing of mouse GL261 GBM cells (Fig. 2f,g). Similarly, MLT-748 treatment enhances the ability CD14-derived human macrophages to kill human GBM cells (Fig. S4c). As controls, we show that MALT1 blockade has no effect on GBM cell apoptosis when the cancer cells were cultured in isolation (Fig. 2h and Fig. S4d). This data support the notion that MALT1 protease inhibition promotes GBM tumor cell-associated macrophages to adopt an active anti-tumor phenotype.

To build upon these observations, we also compared the effect of GBM co-culture on primary murine macrophages isolated from wild-type, MALT1 knockout (MALT1-KO) or MALT1 protease-dead (MALT1-PD) mice (Fig. 3a). MALT1-PD mice harbor a point mutation of the catalytic cysteine within the MALT1 proteolytic domain and are therefore devoid of all MALT1 proteolytic activity^21^. As expected, primary murine WT macrophages polarize toward an M2-like phenotype after co-culture with GL261 murine GBM cells. In contrast, MALT1-PD and MALT1-KO macrophages show a drastic reduction in their capacity to polarize towards a tumor-supportive, M2-like phenotype in response to co-culture with GL261 GBM cells (Fig. 3b,c, left). Instead, MALT1-PD macrophages assume a stronger M1-like phenotype as demonstrated by an increase in CD86 and MHC class II expression on CD11b^+^ cells (Fig. 3b, right). Further, MALT1- PD macrophages demonstrate an increased capacity for tumor cell killing in comparison to WT or MALT1-KO macrophages (Fig. 3d). As a control, we demonstrate that no differences are observed in comparing the survival of WT, MALT1-PD or MALT1-KO macrophages cultured alone (Fig. 3e). Together, these data indicate that inhibiting MALT1 protease activity, specifically within macrophages, promotes their anti-tumor activation status and capacity for killing neighboring malignant glioma tumor cells.

**Fig. 3.**
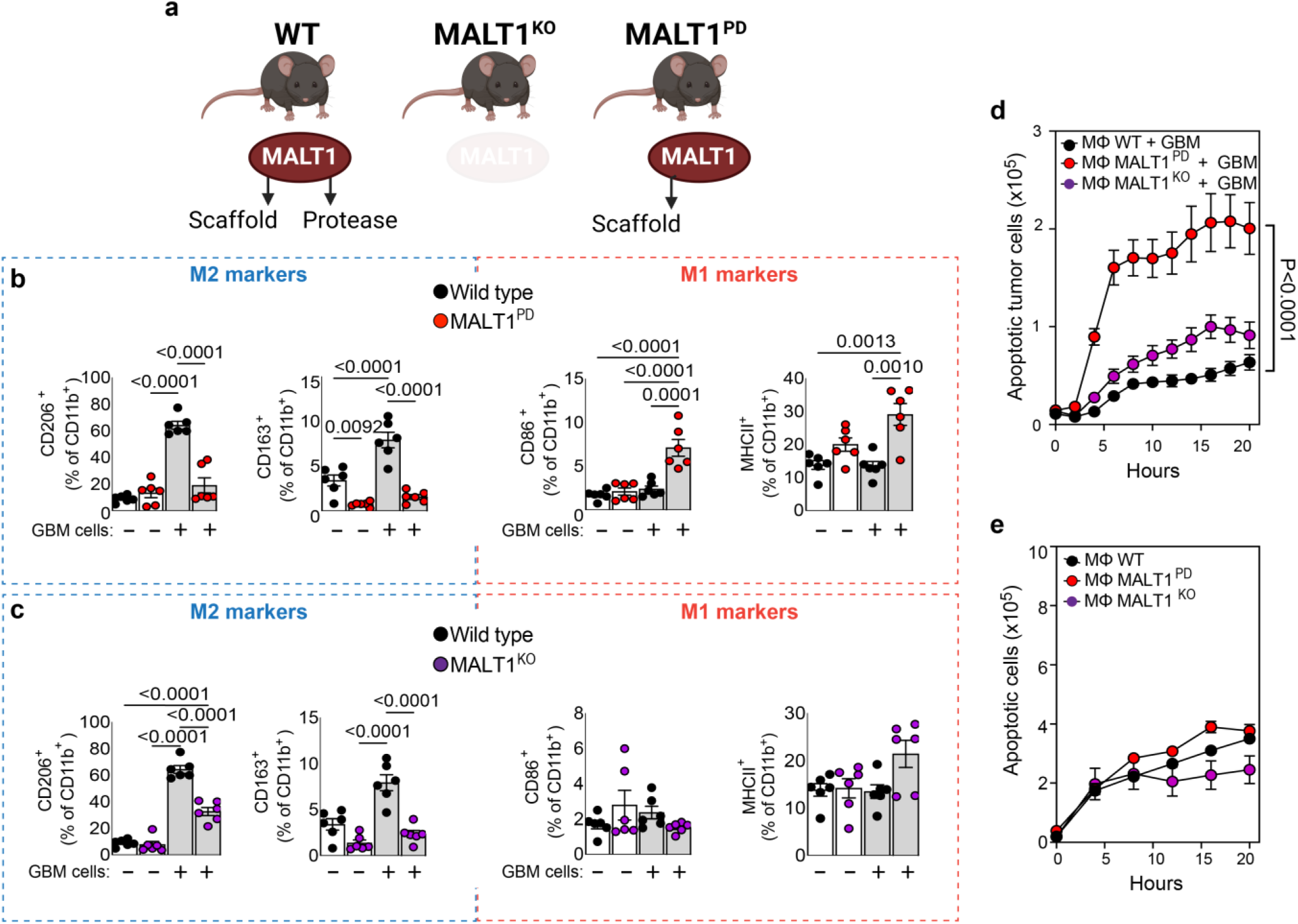
MALT1 genetic targeting reprograms M2-like GBM-associated macrophages into M1-like macrophages. **(a)** Schematic illustrating the WT, MALT1-KO or MALT1-PD mouse phenotype. **(b,c)** Changes in mouse primary macrophage M2/M1-like polarization markers after co-culture with GBM cells. Macrophages were isolated from wild type (WT), MALT1-PD (b) or MALT1-KO (c) mice and co-cultured with GL261 mouse GBM cells. Data are representative of three independent experiments with two technical replicates. **(d)** Quantification from the IncuCyte® live-cell analysis system used to measure macrophage-dependent GBM tumor cell killing. Apoptotic GL261 GBM tumor cells were quantified after co-culture with either WT, MALT1-KO or MALT1-PD macrophages. **(e)** Controls demonstrate that MALT1 status (WT, KO or PD) has no effect on the apoptosis of primary macrophages when they are cultured alone. All values are represented as mean ± SD. Data were analyzed by 1-way ANOVA, followed by Tukey’s multiple-comparisons. P values are indicated in the figure.

Overall, our findings suggest that GBM cancer cells induce the polarization of macrophages towards an M2-like immunosuppressive phenotype and that CBM inhibition, via targeting of either CARD9 or MALT1, reverses this effect, skewing GBM TAMs toward an M1- like phenotype and enhancing their capacity to kill GBM tumor cells. Notably, this M1-like phenotypic skewing requires the presence of GBM cancer cells, since treating either RAW264.7 or human primary macrophages with MLT-748 or BRD5529 in isolation (i.e., in the absence of GBM cells) does not induce this same skewing. The effects of MLT-748 or BRD5529 treatment on the co-culture are likely the result of inhibiting the CBM complex directly within macrophages, and not within GBM cells, because CARD9 is not expressed in GBM cells. Instead of CARD9, the CBM complex within GBM cells contains CARMA3. Therefore, CARD9 inhibition is expected to block CBM activation only in the macrophages and not in the GBM tumor cells. Moreover, the effects of MLT-748 on the cocultures are mimicked when GBM cells, which harbor WT MALT1, are co-cultured with primary MALT1-PD macrophages, which lack MALT1 protease activity.

### Genetic inactivation of MALT1 protease in the TME reduces GBM tumor growth *in vivo* and drives GBM-associated TAMs to assume an anti-tumor M1-like phenotype

We next evaluated the impact of MALT1-protease activity within the TME during GBM progression *in vivo* in mice. For these studies, we generated brain tumors through orthotopic implantation of syngeneic GBM tumor cells into immunocompetent mice, so as to recapitulate key features of the human GBM TME as closely as possible^42^. We evaluated two mouse GBM cell models, GL261 and CT2a, implanted into wildtype (WT) versus MALT1-PD mice. Twenty-one days following implantation, we found that GBM cells produced significantly smaller tumors in MALT1-PD mice than in WT mice (Fig. 4a,b). Additionally, on day 20 post implantation and prior to the time mice in either group had expired, we found that MALT1-PD mice harboring GBM GL261 tumors demonstrated superior locomotor abilities and were more active than WT mice harboring GBM tumors (Fig. S5a-h), suggesting lessened neurological impairment in the MALT1- PD mice. Further, MALT1-PD mice implanted with GL261 GBM cells exhibited a ∼20-day increase in survival relative to WT mice, indicating a survival benefit associated with loss of MALT1 protease activity in the TME. (Fig. 4c). These findings suggest that factors present within the WT TME but absent within the MALT1-PD TME may contribute to tumor progression. Using flow cytometry, we found that tumors from MALT1-PD mice contained a higher overall proportion of CD45^+^ immune cells relative to cancer cells than did tumors harvested from WT mice (Fig. 4d). TAMs (characterized by F4/80 and CD11b expression) made up a greater proportion of the CD45^+^ immune cells present in tumors from WT mice as compared to tumors from MALT1-PD mice (Fig. 4e). Further, of the total TAM population, tumors from WT mice showed a higher fraction with an M2-like immunosuppressive phenotype (CD206^+^Arginase1^+^) while tumors from MALT1-PD mice showed a higher fraction with an M1-like phenotype (iNos^+^MHCII^+^) (Fig. 4e). In addition, the percentage of cells with a monocytic MDSC phenotype (CD11b^+^Ly6C+Ly6G-) within the CD45^+^ cell population was lower in tumors from MALT1-PD mice in comparison to those from WT mice (Fig. 4f). Also, N1-like neutrophils (iNos^+^Arg1^-^), which exhibit increased tumor cell cytotoxicity and immunoreactivity in glioma^43^, made up a greater portion of the neutrophils present in tumors from MALT1 PD-mice in comparison to tumors from WT mice (Fig. 4g).

**Fig. 4.**
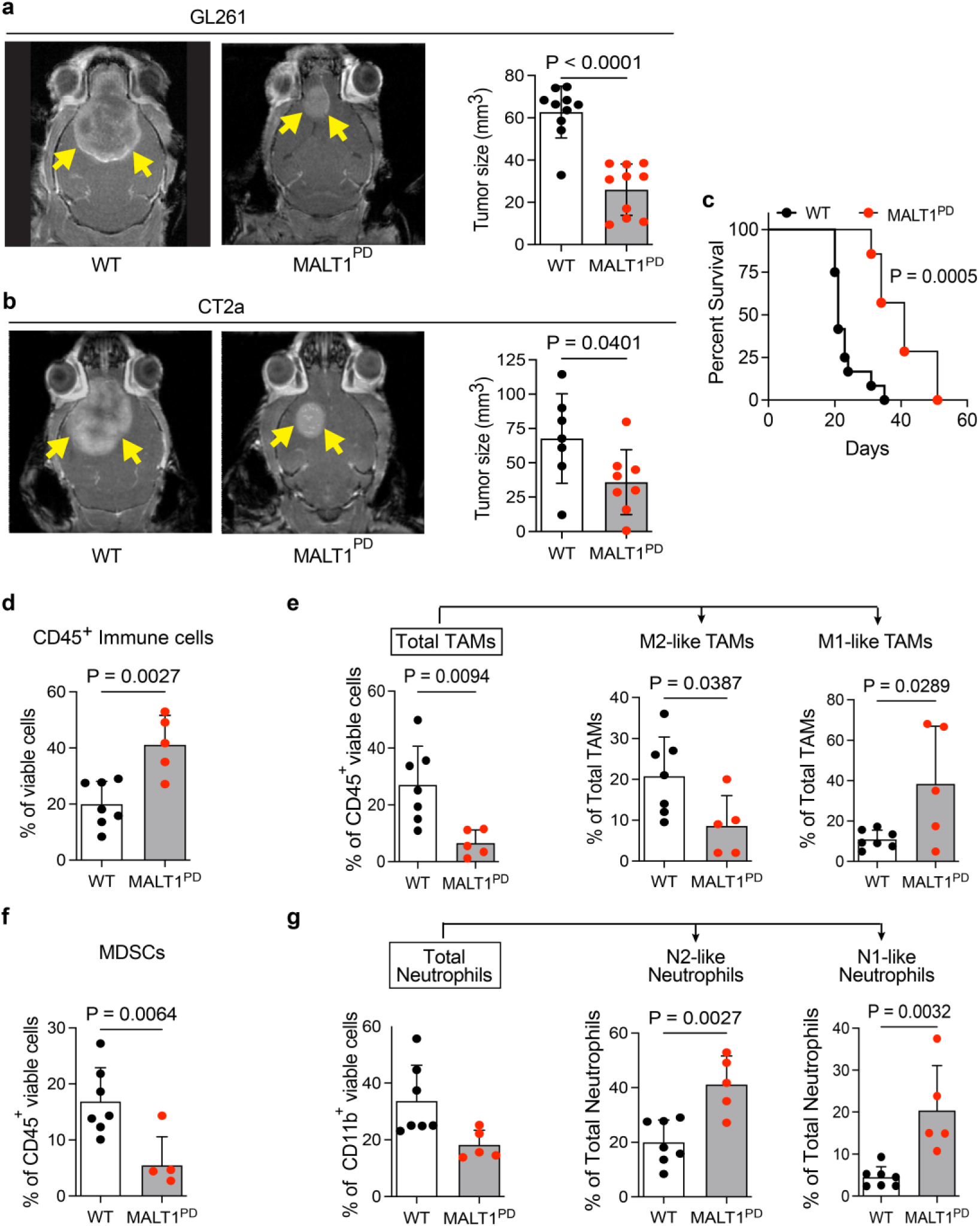
Lack of MALT1-protease activity in the host TME results in smaller GBM tumors and an increase in TME immuno-reactivity. Syngeneic mouse GBM cells (GL261 or CT2a) were implanted into either Wild-type (WT) or MALT1 protease dead (MALT1-PD) C57BL6/J mouse brains through intracranial injection to establish GBM tumors on day zero. **(a-b)** Representative MRI images and quantification of tumor volume after 21 days from WT or MALT1-PD GL261 (a) or CT2a (b) tumor-bearing mice. **(c)** Kaplan–Meier survival curve showing that GL261 GBM tumor-bearing MALT1-PD mice demonstrate increased survival in comparison to WT control mice. A log-rank Mantel–Cox test was performed between the groups with WT (n = 12) and PD (n=8) mice per group. **(d-g)** Flow cytometric quantification of myeloid cells within the tumors, showing the (d) percentage of immune cells (CD45^+^); (e) percentage of total TAMs (CD11b^+^F480^+^), M2-like-TAMs (CD11b^+^F480^+^ Arg1^+^CD206^+^) and M1-like-TAMs (CD11b^+^F480^+^iNos^+^MHCII^+^), (f) M-MDSCs (CD11b^+^Ly6C^+^ Ly6G^-^) and (g) total Neutrophils (CD11b^+^Ly6G^+^Ly6C^-^), N2-like Neutrophils (Arg1^+^CD206^+^) and N1-like Neutrophils (iNos^+^Arg1) in CD45^+^ viable cells in GBM GL261 tumors. Data are shown as mean ± SD from two independent experiments. Statistical significance was determined by two-tailed Student’s t-test; significant P values are indicated in the figure. Each dot represents an independent mouse.

We also analyzed and compared lymphocytes within GBM tumors harvested from WT versus MALT1-PD mice and found no significant differences in the proportion of total T cells, T helper cells, T cytotoxic cells or NK cells (Fig. S6). Notably, we did observe that within the cytotoxic T cell and T helper populations, respectively, the proportion of exhausted T cells and Tregs were significantly lower in tumors from MALT1-PD mice, suggesting an overall more active T cell infiltrate (Fig. S6a-b). We also quantified effector Tregs (eTregs), cells that express higher levels of Treg effector molecules including cytotoxic T cell antigen 4 (CTLA4), which likely contribute to enhanced suppressive activity^44^. We found that the proportion of eTregs are also significantly lower in the MALT1-PD TME (Fig. S6b), again suggesting that the tumor infiltrating T cell population is polarized towards more active anti-tumor immunity in host mice that lack MALT1 protease activity. Together, these findings suggest that MALT1 protease activity within cells of the TME plays a key role in shaping a pro-tumorigenic TME.

### MALT1 protease deficient macrophages in the GBM TME demonstrate reduced immuno- suppressive features and impaired chemotaxis

We next tested for cell-type-specific differences in GBM tumors harvested from WT versus MALT1-PD mice using scRNAseq. Uniform manifold approximation and projection (UMAP) dimension reduction was performed on 21,411 and 21,301 cells, respectively, revealing 26 cell-type clusters in GL261 tumor samples from both WT and MALT1-PD mice (Fig. 5a). Cluster marker genes were consistent with published cell type-enriched genes, including bone marrow derived macrophage (BMDM) clusters (#s 1, 7, 8, 10, 16, 21 and 25) (*LY6C2*, *LY6C1*, *CD14*), the microglia cluster (#4) (*P2RY12*, *TREM2 CSF1R*), the NK cell cluster (#13) (*NKG7*), neutrophil clusters (#13 and 20) (*LY6G*, *ELANE, MPO*, *LCN2*) and lymphocyte clusters (#3 and 19) (*CD3D*, *CD3G*, *CD3E*)^45^ (Fig. S7a).

**Fig. 5.**
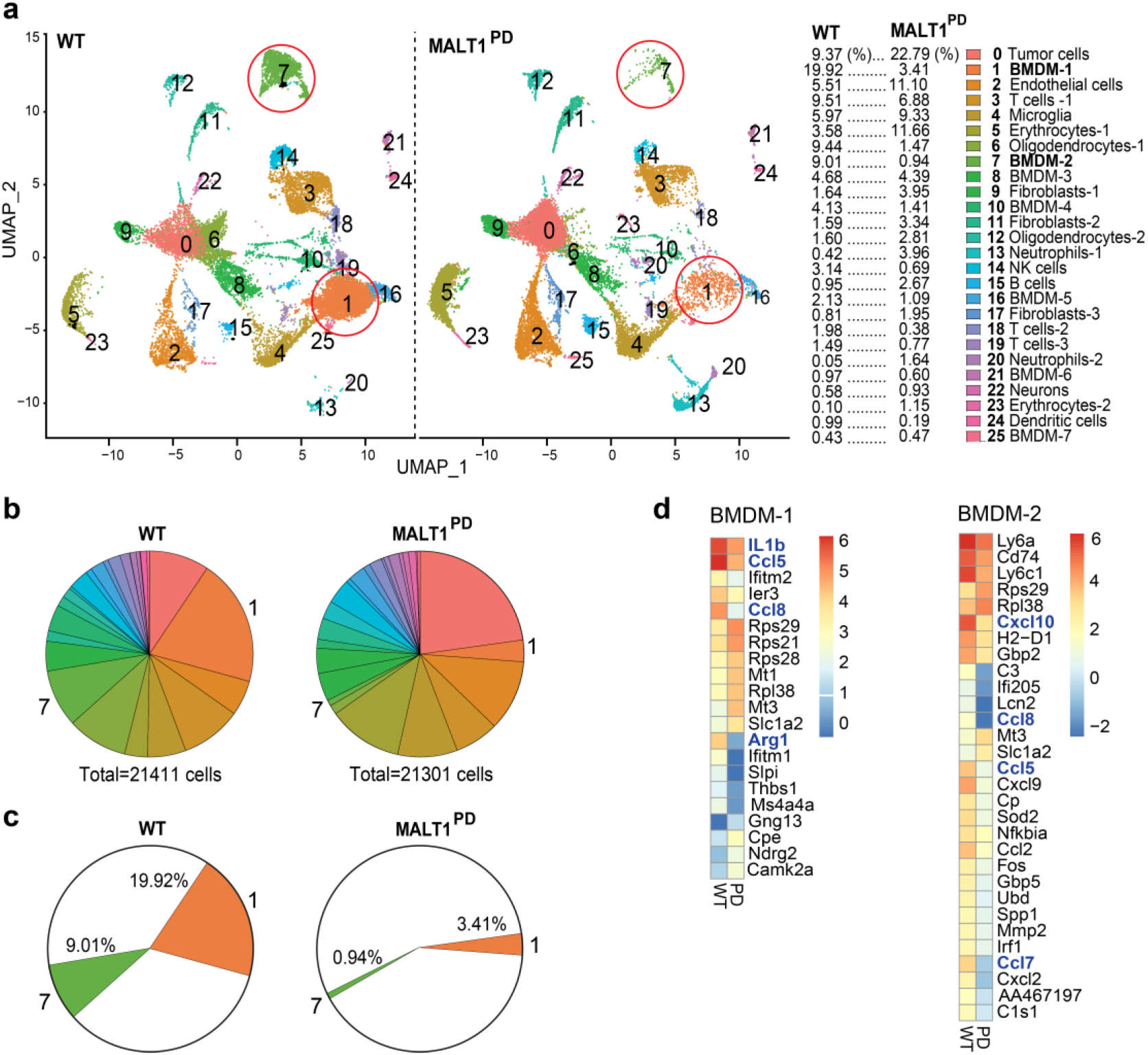
Systemic MALT1-protease deficiency is associated with decreased macrophage infiltration and reduced macrophage immunosuppressive polarization in the GBM-TME. **(a)** GL261 GBM tumors which grew after implantation into Wild Type (WT) or MALT1-PD mice (n = 3 per group) were dissected, and viable cells were subjected to transcriptome sequencing. The UMAP plot of 26 clusters is shown. Cluster 1 (BMDM-1) and cluster 7 (BMDM-2) are highlighted with red circles. **(b)** Annotated clusters of intratumoral cells are represented by pie charts. Relative cell numbers for each cluster are listed. Pie chart colors match the cluster colors used in the UMAP plots. **(c)** Pie charts highlighting the most prominent differences (BMDM-1 and BMDM-2; clusters #1 and #7) between the GBM TME of MALT1-WT and MALT1-PD mice. **(d)** Heatmap representation of the top differentially expressed genes in the BMDM-1 and BMDM-2 macrophage clusters from GBM-bearing WT vs MALT1-PD mice. Heatmaps reflect log2 average expression of top differentially expressed genes.

The scRNAseq analysis revealed that MALT1 protease deficiency in host mice is associated with decreased infiltration of bone-marrow derived macrophages (BMDMs) into the TME of GL261 GBM tumors (41% of total cells in WT group vs 12% in MALT1-PD group) (Fig. 5a,b) This was primarily due to a reduction of the two major BMDM clusters (BMDM-1 and BMDM-2) which were significantly more prevalent in the tumors in WT mice (19.92% in WT vs 3.41% in MALT1-PD; and 9.01% in WT vs 0.94% in MALT1-PD group, respectively) (Fig. 5c). These results are in agreement with our flow cytometry data which shows reduced TAMs in tumors from MALT1-PD mice in comparison to tumors in MALT1-WT-mice (27% of CD45^+^ cells in the WT group vs 6.5% in the MALT1-PD group) (Fig. 4e). Notably, the impact on TAM populations was restricted to BMDMs, as we did not observe a decrease in the microglial population (cluster #4) in tumors from MALT1-PD mice versus WT mice (Fig. 5a,b).

In addition to revealing this reduction in BMDMs, differential gene expression analysis demonstrated that the GBM-associated BMDMs harvested from MALT1-PD mice show decreased expression of genes that are characteristic of M2-polarization^46^. Within the BMDM-1 cluster (the dominant cluster present in the TME of tumors in WT mice), this included decreases in *ARG1*, *CCL8*, and *CCL5* (Fig. 5d). In addition, we noted that GBM-associated BMDMs from MALT1- PD mice showed decreased expression of selected chemokines and cytokines, many of which are known for their pro-tumorigenic effects^47,48^. These included *CCL5, CCL8,* and *IL1B* within the BMDM-1 cluster, as well as *CXCL10, CCL8, CCL5,* and *CCL7* in the BMDM-2 cluster (Fig. 5d). The other BMDM clusters from GBMs in MALT1-PD mice showed similar downregulation of these and other secreted factors (Fig. S7b). Overall, the scRNAseq data suggest that bone marrow derived macrophages from MALT1-PD mice have reduced capacity to infiltrate into the GBM microenvironment. Once there, these macrophages may also be functionally impaired, unable to adopt an M2-phenotype and therefore less effective at establishing and maintaining an immunosuppressive microenvironment.

Because we noted that MALT1-protease deficiency in host mice is associated with a significant decrease in the infiltration of bone marrow derived macrophages into the GBM microenvironment (Fig. 4,5), we decided to investigate the impact of inhibiting the CBM complex/MALT1 on macrophage migration and chemotaxis. Specifically, we used two complementary approaches to directly test how CBM/MALT1 blockade affects the ability of macrophages to migrate towards brain tumor cells. First, we performed a scratch wound assay in which we treated RAW267.4 murine macrophages with GL261 GBM tumor cell conditioned media (CM), and then compared their migration -/+ treatment with MALT1 or CARD9 inhibitors (see schematic, Fig. 6a). We first demonstrated that GL261 CM effectively induces migration of macrophages in this system (Fig. 6b). We then found that treatment of macrophages with either of the MALT1-protease inhibitors, mepazine or MLT-748, abrogates this CM-induced macrophage cell migration in a dose dependent fashion (Fig. 6c,d). A similar trend toward decreased migration was observed with the CARD9 inhibitor, BRD5529 (Fig. 6e). Second, we utilized a transwell system in which macrophages were cultured in the top chamber and GL261 GBM tumor cells in the bottom chamber. We then compared migration of macrophages toward GBM cells, -/+ MALT1 protease blockade or CARD9 inhibition (see schematic, Fig. 6f). As a control, we first demonstrated that macrophages migrate toward the GBM cells in this system (Fig. 6g). We then found that both MALT1-protease inhibitors as well as the CARD9 inhibitor, BRD5529, block the ability of macrophages to migrate towards the GBM tumor cells (Fig. 6h). Finally, we also found that macrophages isolated from MALT1-PD or MALT1-KO mice show strikingly impaired chemotaxis in comparison to WT macrophages (Fig. 6i). These findings, together with our flow cytometry and scRNAseq analyses of GBM tumors, support the notion that MALT1 activity within macrophages plays a key role in promoting macrophage migration into the GBM tumor microenvironment.

**Fig 6.**
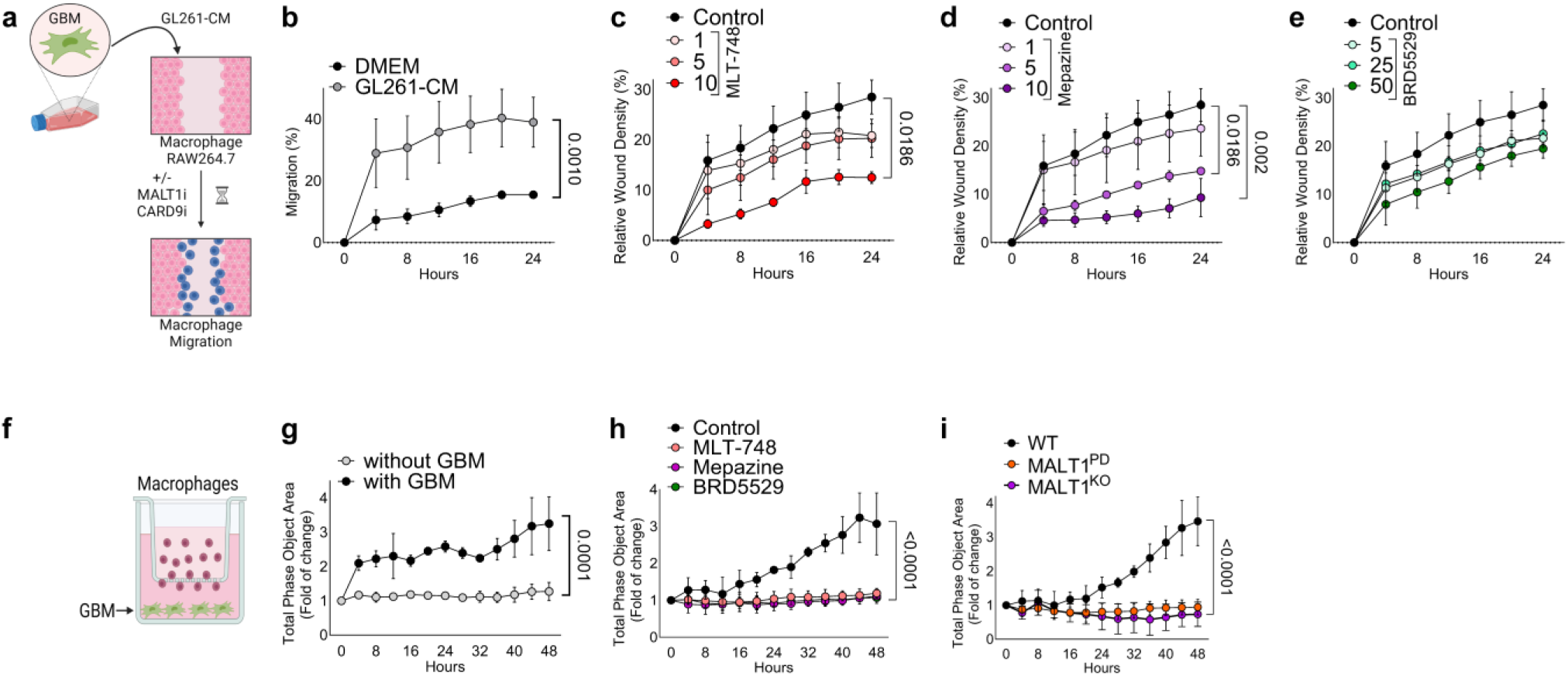
CBM complex activity promotes GBM-associated macrophage migration. (**a**) Schematic of macrophage cell migration assay. RAW264.7 macrophage cell migration was monitored in a real-time scratch assay, using the IncuCyte system. Blue colored cells are meant to represent macrophages that have migrated into the scratch wound. **(b)** Conditioned media (CM) from GL261 GBM cells enhances migration of macrophages in comparison to DMEM control. (**c-e**) Macrophages were exposed to GL261-GBM CM and treated with increasing concentrations of MLT-748 (c), mepazine (d) or BRD5529 (e). Control cells were exposed to GL261-CM with equivalent volumes of DMSO. Wound density (closure) is plotted as a continuous function of time. (**f)** Schematic of transwell migration assay. **(g)** GL261 GBM tumor cells induce the migration of primary macrophages isolated from WT mice. **(h)** MALT1 protease inhibitors (MLT-748 or mepazine) and CARD9 inhibitor (BRD5529) each prevent GBM cell-induced macrophage transwell migration. **(i)** Primary macrophages from WT, MALT1-PD or MALT1-KO were tested for chemotactic migration towards GL261-GBM cells in the bottom chamber. Data shown is from three independent experiments. The P values were calculated by One-way ANOVA.

### Pharmacologic MALT1 protease inhibition increases immuno-reactivity of GBM TAMs, reduces GBM tumor progression and enhances treatment response when combined with standard-of-care chemotherapy

The reduction in M2-like TAMs together with the improved overall survival we observed in the MALT1-PD mice harboring GBM tumors prompted us to consider whether pharmacologic inhibition of MALT1 protease activity could similarly reduce tumor immune suppression and drive anti-tumor immune activity within the GBM TME. To complement our genetic approach to blocking MALT1 proteolytic activity in the TME, as shown in Fig. 4, we next evaluated the effect of treating mice harboring GBM tumors with the BBB-penetrant, allosteric MALT1 protease inhibitor, mepazine, a commercially available phenothiazine derivative^49^. Specifically, starting on day 5 after orthotopic implantation of GL261 GBM cells, C57BL/6 mice were treated with vehicle, standard-of-care chemotherapeutic agent, Temozolomide (TMZ)^50^, mepazine, or the combination of TMZ + mepazine (see schematic, Fig. 7a).

**Fig 7.**
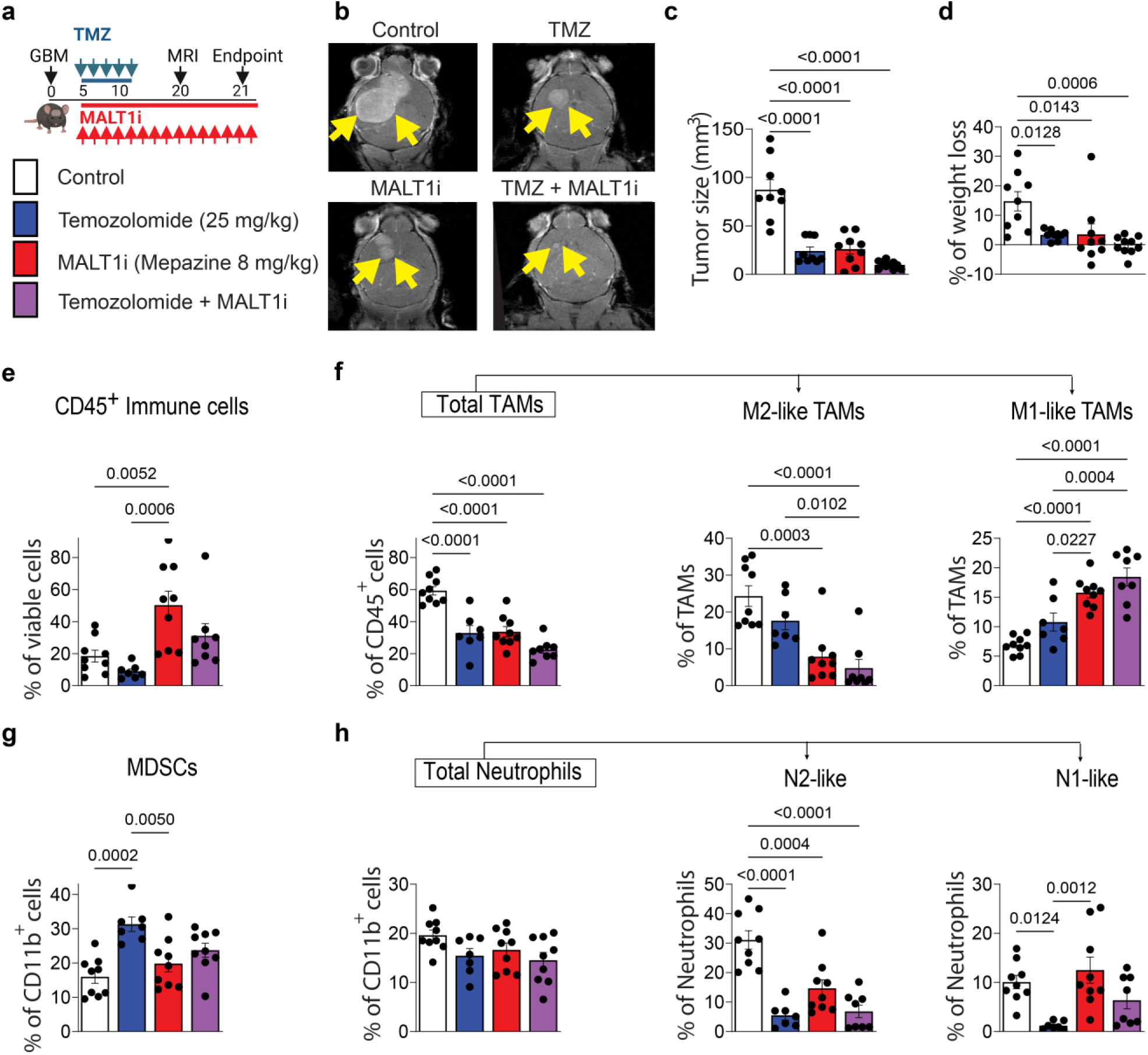
Pharmacologic MALT1 protease inhibition results in reduced GBM progression, increased immuno-reactivity of GBM TAMs, and enhanced response to standard-of-care chemotherapy. **(a)** Schematic illustrating the Mepazine (MALT1i) and Temozolomide (TMZ) treatment schedule. Syngeneic mouse GL261 cells were transplanted into mouse brains through intracranial injection to establish GBM tumors. Five days after transplantation, tumor-bearing mice were treated with vehicle control (DMSO 5%), TMZ (25 mg/kg IP QD for 5 days), MALT1 inhibitor (MALT1i) Mepazine (8 mg/kg IP BID for 16 days), or TMZ + Mepazine. MRI was performed on day 20 to compare tumor growth. Mouse brains bearing GBM tumors were collected for further analyses on day 21. **(b)** Representative MRI images and **(c)** quantification of tumor volume from GBM tumor-bearing mice. **(d)** % of weight loss on day 21. **(e-h)** Flow cytometric quantification of GBM TME immune composition showing the **(e)** percentage of immune cells (CD45^+^), **(f)** proportion of TAMs (CD11b^+^F480^+^), M2-like-TAMs (CD11b^+^F480^+^CD206^+^Arg1^+^) and M1-like-TAMs (CD11b^+^F480^+^iNos^+^MHCII^+^), **(g)** M-MDSCs (CD11b^+^Ly6C^+^Ly6G^-^) and **(h)** total neutrophils (CD11b^+^Ly6G^+^Ly6C^-^), N2-like neutrophils (Arg1^+^CD206^+^) and N1-like neutrophils (iNos^+^Arg1^-^). Data are shown as mean ± SD. n = 9 for control, n=9 for TMZ, n=9 for MALT1i and n=10 TMZ + MALT1i per group. Statistical significance was determined by one way ANOVA followed by Tukey; significant P values are indicated in the figure.

Systemic immunosuppression is a hallmark feature of GBM which is typically seen even before initiation of treatment^51^. Furthermore, TMZ, the most commonly used agent for the treatment of GBM, is well known to induce lymphopenia and further exacerbate systemic immunosuppression^50^. In light of these effects of GBM and of TMZ, we decided to first investigate the systemic effects of our treatment arms by using flow cytometry to characterize the immune cell populations in the spleens of GBM-bearing mice (Fig. S8). Treatment with mepazine alone had no major effects on the appearance of spleens or on most immune cell populations isolated from spleens (Fig. S8a-i), with one exception being that mepazine treatment resulted in a decrease in splenic MDSCs (Fig. S8d). The addition of mepazine to TMZ had no effects on the total number of macrophages, neutrophils, lymphocytes, or NK cells in the spleen but did result in a decrease in the relative proportion of M2-like macrophages and N2-like neutrophils in comparison to TMZ alone (Fig. S8c,e). We found that, as expected, treatment with TMZ alone caused a decrease in the percentage of splenic T helper cells and T cytotoxic cells and this effect was unchanged by the addition of mepazine to the TMZ (Fig. S8g). Overall, the results suggest limited effects of mepazine treatment on systemic immune cell populations over the course of this 16-day treatment.

Consistent with our analysis of the MALT1-PD genetic model, MRI imaging demonstrated that monotherapy with the MALT1 protease inhibitor, mepazine, substantially restrains GBM tumor growth *in vivo* (Fig. 7b,c). Results with mepazine treatment alone were similar to results with TMZ treatment alone, while the combination showed a trend toward further growth inhibition (Fig. 7b,c). Furthermore, the addition of MALT1 inhibitor reduces the GBM-induced weight loss seen in the TMZ-treated mice (Fig. 7d).

Immunophenotyping of the GBM TME showed that pharmacologic MALT1 protease inhibition with mepazine mimics the phenotype observed in MALT1-PD mice implanted with GBM tumors (see Fig. 4), at least in part. Similar to the GBM TME in MALT1-PD mice, the GBM TME in mepazine-treated mice showed enhanced presence of CD45^+^ immune cells in comparison to control vehicle-treated mice (Fig. 7e). In addition, while TMZ treatment was associated with a trend toward decreased immune infiltration, the addition of mepazine to the TMZ partially reversed this effect (Fig. 7e). Mepazine treatment also resulted in a decrease in M2-like TAMs and an increase in M1-like TAMs (Fig. 7f), similar to what we observed in the MALT1-PD mice (see Fig. 4e). Notably, combination mepazine plus TMZ treatment leads to a greater decrease in M2- like TAMs and a greater increase in M1-like TAMs than TMZ alone (Fig. 7f). While TMZ alone increased the number of MDSCs and decreased N1 neutrophils, we observed a trend suggesting that concomitant treatment with mepazine may partially reverse these effects. In the lymphocyte compartment, both MALT1-PD mice and mepazine-treated mice demonstrated a reduced presence of exhausted T cells within the GBM tumor (Fig. S6a and S9a, respectively). Further, combination therapy of TMZ plus mepazine resulted in a decrease in Effector Tregs and an increase in NK cells in the GBM TME.

In summary, our *in vivo* studies demonstrate that systemic MALT1 protease inhibition, via either pharmacologic treatment or genetic modification of the host, is effective at reducing the immunosuppressive microenvironment directed by malignant GBM cells and established by infiltrating macrophages. As such, MALT1 inhibition alone is effective at abrogating GBM tumor progression. Further, the addition of a MALT1 inhibitor to TMZ treatment may enhance the immunoreactivity of the GBM TME in comparison to TMZ alone and may thereby enhance treatment response to this standard-of-care chemotherapeutic. Together, the findings presented here suggest that the CBM/MALT1 signaling axis in GBM TAMs holds promise as a new therapeutic target for GBM.

## Discussion

Our study reveals a key mechanism by which glioblastoma tumor cells escape immunosurveillance through inducing MALT1 activity in TAMs to create an immunosuppressive environment. Using co-culture studies, we show that GBM tumor cells can induce NF-κB activation within macrophages via a CARD9-MALT1-dependent mechanism. This is associated with enhanced migration and chemotaxis of macrophages towards GBM cells, along with their adoption of an M2-like polarized state. Importantly, pharmacologic blockade of either CARD9 or MALT1 abrogates all of these GBM-driven effects on macrophage phenotype. Further, our genetic studies in mice show that loss of MALT1-protease activity within the host TME results in reduced GBM tumor growth and enhanced mouse survival. Finally, our additional *in vivo* analyses with systemic mepazine administration demonstrate the efficacy and feasibility of pharmacologically targeting MALT1-protease as a new immunomodulatory approach to treating GBM. As a single agent, mepazine drives an *in vivo* M2-like to M1-like TAM phenotype switch within the GBM TME and reduces tumor growth. Notably, our findings suggest that mepazine has no effect on GBM tumor cell survival or proliferation *in vitro*. This combination of findings suggests that the major effects of targeting MALT1 protease *in vivo* with mepazine are due to cancer cell-extrinsic alterations of the TME that hinder tumor growth, rather than a cancer cell-intrinsic effect. Specifically, our work suggests that MALT1 protease inhibition activates an antitumor immune mechanism that reprograms expression of immune-associated genes in macrophages. This work demonstrates the potential for using a MALT1 inhibitor in combination with the standard-of-care chemotherapy, TMZ, since systemic MALT1 inhibition may prevent some of the undesired immunosuppressive effects of TMZ and reduce tumor growth beyond what can be achieved with TMZ alone.

To date, most work investigating approaches for inhibiting the CARD-BCL10-MALT1 signalosome have focused on targeting the effector protein of the complex, MALT1, through the use of MALT1-specific protease inhibitors, rather than on targeting other components of the signalosome. MALT1 possesses two major functions, acting as a scaffolding protein and as a protease^8^. Blocking MALT1-protease activity, without inhibiting MALT1 scaffolding activity, has the potential for abrogating not only the immunosuppressive actions of TAMs in the TME, but also of Tregs, since previous work has shown that selectively blocking MALT1 protease activity disproportionately impacts Tregs while allowing continued T effector function^31^. This observation has added therapeutic implications since MALT1 protease inhibition could allow for enhanced T cell-mediated anti-cancer activity. A potential limitation to this approach, however, could relate to previous reports which have shown that while MALT1-KO mice present with severe immunosuppression because they lack lymphocyte activation due to the loss of both MALT1 scaffold and protease functions, MALT1-PD mice instead develop autoimmunity^18,19,21^. The systemic inflammation associated with MALT1 protease deficiency is thought to be caused by residual immune activation in T effector cells due to preserved MALT1 scaffolding function, which in the absence of MALT1 protease activity, leads to a relative increase in the ratio of T effector to Treg activity and shifts the balance from tolerance to autoimmune activation^31^. While we find that targeting MALT1 protease in a GBM-bearing immunocompetent mouse model for up to 16 days does not result in appreciable inflammatory toxicity, future studies may reveal that the inflammation associated with long term MALT1 protease inhibitor treatment is a limitation of this therapy.

Our study shows that the CARD9-MALT1 signaling axis is critical for the GBM-dependent immunosuppressive polarization of TAMs, suggesting that CARD9 is essential for mediating the TAM response to GBM cells. CARD9 is one of several CARD family members that coordinate CBM complex assembly and MALT1 activation in immune and other cell types^38^. In contrast to MALT1, which is widely expressed in most or all cell types, including T cells, CARD9 is a component of the CARD-BCL10-MALT1 signalosome that is expressed primarily in the myeloid compartment^38^. CARD9 acts to relay signals from multiple receptor subtypes, including Toll-like and C-type lectin receptors, by anchoring the CARD-BCL10-MALT1 signalosome to drive downstream signaling in macrophages^38^. As an alternative to MALT1 protease inhibition, targeting CARD9 within GBM TAMs could represent an approach to reducing TAM-dependent immunosuppression in the GBM TME while simultaneously avoiding the potential for the systemic autoimmune response associated with MALT1 protease inhibition. Our work here with the CARD9 inhibitor, BRD5529, which abrogates activation of the CARD9-BCL10-MALT1 signalosome within macrophages, serves as an initial proof-of-concept for this approach.

Here, we have shown that GBM-associated macrophages are highly affected by the loss of MALT1-protease activity whereas microglia may be less affected. TAMs in GBM are composed of bone marrow–derived macrophages (BMDMs) and brain-resident microglia. The main differences between these two TAM cell types include (1) BMDMs originate from progenitor cells in the bone marrow while microglia originate from the yolk sac, (2) microglia are found to be prominent in the peri-tumoral areas, while macrophages are dispersed within the tumor bulk and are more abundant in the perivascular areas and (3) BMDMs and microglia are characterized by distinct transcriptional and morphological patterns^7^. Our data indicates that the relative number of macrophages within the GBM TME is significantly reduced in MALT1-PD mice as compared to WT mice. This decrease is not observed for microglia within the TME. These findings could reflect the fact that MALT1 protease activity plays a key role in macrophage recruitment into the TME, whereas resident microglia are present within the brain and are not recruited into the tumor from the periphery. Further studies are needed to characterize the TAM spatial distribution within GBM tumors and to elucidate how/why GBM-associated microglia respond differently to loss of MALT1 protease activity than do BMDMs.

One limitation of this current study is that pharmacologic MALT1 protease inhibitor treatment was conducted in murine GBM models that do not fully capture the diverse heterogeneity of human gliomas. One potential approach to further informing the translation of our findings to clinical treatment in human patients with GBM would be to evaluate MALT1 protease inhibition in immune-humanized patient-derived GBM xenograft models and/or patient- derived GBM organoids. It may also be informative to evaluate additional MALT1 protease inhibitor compounds in future preclinical studies. Our team chose to use mepazine as the pharmacologic agent to target MALT1 protease in GBM *in vivo*. We focused on this agent because thus far, mepazine is the only MALT1 protease inhibitor known to demonstrate BBB-penetrance^52^. Mepazine and other related phenothiazines have been used clinically as antipsychotic drugs and second-generation derivatives of mepazine have recently become available^53^. One potential next step for our analysis would be to evaluate MPT-0118, an S-enantiomer of mepazine that is currently being tested in a Phase 1 clinical trials in patients with advanced or metastatic refractory solid tumors (NCT04859777). This S-enantiomer possesses an approximately 9-fold higher potency than the corresponding (R)-isomer^53^.

There are major clinical implications of this study. Our results identify the CARD9- containing CBM signalosome as a potential target for the development of new immunotherapeutic strategies for brain tumors. GBM, recognized as the most aggressive primary brain tumor, poses significant treatment challenges due to its rapid growth, capacity to infiltrate brain tissue, molecular diversity, and resistance to treatment^1^. Despite two decades of relentless pursuit of novel therapeutic approaches for GBM, there has been only limited progress in improving patient outcomes. Numerous obstacles impede the effective application of immune-modulating agents for the treatment of GBM, including the presence of a large population of immunosuppressive myeloid cells, low numbers of tumor-infiltrating lymphocytes and other immune effector cell types, the blood-brain barrier, and extensive tumor heterogeneity^4^. In our preclinical mouse studies, the magnitude of the antitumor effect conferred by treatment with the MALT1 protease inhibitor mepazine is similar to the effect seen with administration of TMZ, the main chemotherapeutic agent that has been utilized for the treatment of GBM during the past 20 years. It is important to highlight that since the initial discovery of MALT1 protease activity in 2008, there has been an increasing interest in the development of MALT1 protease inhibitors suitable for clinical use^53^. MALT1 protease inhibitors are already being tested in clinical trials in human patients, indicating the potential for rapid translation of our findings.

The current standard-of care treatment paradigm for GBM, known as the Stupp regimen, entails maximal surgical tumor resection followed by a combination of radiotherapy and chemotherapy^54^. Despite this aggressive approach, almost all patients suffer from tumor recurrence after receiving standard treatment and TMZ increases two-year survival rates from 10% to only 25% in glioma patients, and five-year survival rates, from insignificant to only 10%. TMZ has been shown to cause lymphopenia, to increase the proportion of Tregs in the TME, and to potentially enhance immunosuppression in the myeloid compartment^50^. These immunomodulatory effects of TMZ have been shown to negatively impact GBM response to various immunotherapeutic strategies^55^. Our analysis suggests that the combination of MALT1 inhibitor treatment plus TMZ chemotherapy is feasible and may enhance antitumor immunity in GBM. Our results also suggest that targeting MALT1 may potentially be effective both as monotherapy and as combination therapy with TMZ. Resistance to TMZ is one of the significant limitations in the treatment of GBM. In this context, the immune TME is thought to be critically involved in resistance to TMZ-therapy because chemoresistant GBM cells promote M2-like polarization of macrophage to a greater degree than chemosensitive cells, which sustains tumor growth^56^. Future studies could be designed to evaluate if the effects of MALT1 protease inhibition on the GBM TME would be beneficial in the context of TMZ chemoresistance.

In summary, our work establishes that MALT1 protease inhibition reverses GBM tumor cell-induced, macrophage-dependent immunosuppression. The efficacy of both genetic and pharmaceutical inhibition of MALT1 protease in reducing tumor growth and reprogramming the TME in GBM mouse models strongly supports the further development of MALT1 inhibitor-based immunotherapeutic strategies for brain tumors. MALT1 protease inhibitors may provide a key opportunity to overcome immunosuppression in tumors with high macrophage infiltration.

### Methods Study design

The purpose of this study was to evaluate the role of MALT1 in the glioblastoma TME and to investigate the underlying molecular mechanisms by which MALT1 influences immune cell actions in this setting. Specific CARD9 and MALT1 inhibitors were used to assess the therapeutic potential of targeting the CBM complex using both *in vitro* and *in vivo* analyses. *In vitro* studies were performed using Western blotting, flow cytometry, reporter assays, colony formation assays, cell proliferation assays, migration, and invasion assays. For in vivo studies, multiple mouse models were utilized. GBM tumor cells were implanted into the brains of MALT1-PD or WT mice and the resultant tumor sizes were monitored and compared using MRI. Tumors were harvested, and their immune compositions were analyzed and compared by flow cytometry and single-cell RNA-seq. In a complimentary analysis, mice bearing GBM tumors were treated with/without MALT1 protease inhibitor alone or in combination with standard-of-care chemotherapy. The composition of the GBM TME and the resulting GBM tumor growth within each treatment arms were compared. All experiments were randomized and blinded where possible. Sample sizes were determined based on expected effect sizes from pilot experiments. In general, group sizes of five or more mice were used. Differences in tumor growth were tested using one-way analyses of variance (ANOVAs) or T test.

### Animals

All experimental animal procedures were approved by the Institutional Animal Care and Research Advisory Committee of University of Pittsburgh (22071589). Mice were maintained under pathogen-free and temperature- and humidity-controlled conditions with a 12-h light/12-h dark cycle. C57BL6/J mice were purchased from Jackson Lab. All mice used for tumor experiments were male and females between 4 and 8 weeks old, with treatment arms matched for age and sex.

### Cell lines

The mouse GL261 and CT2a and the human U87MG cell lines were kindly provided by Baoli Hu (University of Pittsburgh) and cultured in DMEM (Gibco) supplemented with 10% of fetal bovine serum (FBS; Gibco). Mouse Macrophage (RAW264.7) was obtained from the American Type Culture collection (ATCC) and cultured in DMEM 10% FBS. RAW-Blue™ cells were from InVivoGen and grown in DMEM with 10% heat-inactivated FBS, 100 μg/ml Normocin and Zeocin. All cells were grown at 37C in a 5% CO2 incubator. Cell lines were regularly monitored for mycoplasma contamination using the mycoplasma MycoAlert Detection Kit (catalog no.: LT07–318, Lonza).

### CD14 derived primary human macrophages

Human blood was obtained from deidentified donors from Vitalant under Institutional Review Board (IRB) #21080109, which has a "not human research classification". Peripheral blood mononuclear cells (PBMCs) were isolated by Ficoll Paque Plus (catalog no.: 17144002, Cytiva) by gradient centrifugation as previously described^41^. PBMCs were used for the isolation of CD14^+^ cells by negative selection using AutoMACS with the cell isolation kit (catalog no.: 130-096-537, Miltenyi). Purified monocytes (<90%) were cultured in RPMI + 10% FBS + 50 ng/ml Recombinant Human GM-CSF (PeproTech, catalog no.: 300-03) for 6 days to induce macrophage differentiation.

### Preparation of mouse-derived macrophages

Murine macrophages were prepared and cultured according to an established protocol^56^. Briefly, macrophages were collected by lavage of the murine peritoneal cavity with PBS/3% FBS medium. Cells were centrifuged and suspended in DMEM/FBS-free medium. Macrophages were allowed to attach for 30 min. Then, medium containing unattached cells was removed, and cells were grown in DMEM 10% FBS.

### In vitro investigations of MALT1 in GBM cancer cells

#### MALT1 Knockdown

ON-TARGET plus SMART pool siRNAs targeting MALT1 (catalog no.: LQ-005936-00-0020) were obtained from GE Dharmacon. Nontargeting siRNA pools (catalog no.: D-001810–10–50) were used as controls. Lipofectamine RNAiMAX (catalog no.: 13778150, Thermo Fisher Scientific) was utilized to reverse transfect SMARTpool siRNAs (20 nmol) into cells following the manufacturer’s protocol. The knockdown efficiencies for MALT1 were determined by immunoblot assays.

### GBM Cell migration

GL261 and U87MG glioma cell lines were seeded in 96-well plates (5 × 10^4^ cells/well). Cells were exposed to MALT1-siRNA complexes or treated with MLT-748 (5 μM) in DMEM/10% FBS.

After 72 h, a cell monolayer wound was created in reduced FBS conditions (DMEM/0.5% FBS). Next, DMEM/0.5% FBS medium was replaced with fresh reduced FBS medium in the presence of MALT1-siRNA complexes or MLT-748. Control cells were exposed to DMEM/0.5% FBS containing DMSO or Nontargeting siRNA pools. 2D cell migration assays were performed following the Incucyte ZOOM 96-well Scratch Wound Cell Migration assay protocol (Sartorius).

### GBM Cell viability

5×10^3^ cells/well were seeded in 96-well plates in the absence or presence of increasing drug concentrations (0-20 μM) or MALT1-siRNA-lipofectamine complexes. Following 72 h of treatment, cell viability was assessed by 3(4,5-dimethyl)-2,5diphenyltetrazolium bromide (MTT) assay (catalog no.: M5655, Sigma Aldrich). MTT solution was added to the incubation medium in the wells at a final concentration of 0.5 mg/mL. The cells were left for 90 min. The medium was then removed, and plates were shaken with 50µl of DMSO. Optical density of each well was measured at 570 nm. Results were expressed as percentage of control according to the following formula: cell viability rate (%) = (OD570 of treated cells/OD570 of control) × 100%.

### GBM cell Clonogenic Assay

Cells were exposed to MLT-748 (1, 5 and 10 μM) or MALT1-siRNA. After 72 h, the treatment was removed; cells were washed twice with PBS, harvested, and seeded at a density of 100 cells/well in a 6-well plate. After 14 days, colonies were fixed with 100% methanol, followed by staining with 0.1% crystal violet. The number of colonies was counted, and colonies were photographed for analysis. Data were expressed as the percentage of colony number compared with DMEM/10% FBS or Control-siRNA treated cells.

### In vitro investigations of MALT1 in macrophages

#### Macrophage phenotype

Control macrophages were maintained in DMEM with 10% FBS. GBM cells were co-cultured with macrophages for 48 h. Macrophages were phenotype by flow cytometry. In experiments with the MALT1 inhibitor, macrophages were exposed to MLT-748 (5 µM, catalog no.: HY-115466, Med Chem Express) or BRD5529 (50 µM, catalog no.: HY-115497) for 2 hours prior to addition of tumor cells and during the entire co-culture time. To study polarization, changes in expression of M1/M2 markers on macrophages were measured by flow cytometry using a Fortessa (BD Biosciences) cytometer after surface staining with specific antibodies (Supplementary Table 1).

### In vitro cytotoxicity assay

Macrophages (effector cells) were co-incubated with GBM tumor cells (target cells) at a ratio of 1:2 (1×10^4^ GBM Cell Tracker red marked tumor cells per well). Target cell killing was assayed by Caspase-3/7 Dye for Apoptosis and monitored over time using Incucyte.

### MALT1-NF-κB activation

RAW 264.7-GloSensor cells were generated by lentiviral infection following antibiotic selection. The assay was performed according to the manufacturer’s instructions. Briefly, 2 × 10^4^ RAW-blue cells or RAW 264.7-GloSensor cells were incubated with 2 × 10^4^ GBM cells alone or in combination with a MALT1 inhibitor (MLT-748) or a CARD9 inhibitor (BRD5528) for 48 hours under standard conditions. After incubation, 20 µL of media was collected and incubated with 200 µL QUANTI-blue reagent (cat no.: rep-qbs, Invivogen) and optical density was measured at 655 nm using TECAN spectrophotometer (TECAN) for NF-kB activity. For MALT1 activity, Luciferase Assay Reagent II (cat no.: E1910, Promega) was added to the cell lysate and luminescence was measured.

### Immunoblot Assay

Cell lysates were prepared with RIPA buffer (catalog no.: 89901, Thermo Fisher Scientific) containing HALT Protease and Phosphatase Inhibitor cocktail (catalog no.: 78440, Thermo Fisher Scientific), loaded in 4% to 15% Mini-PROTEAN TGX Precast Protein Gels (catalog no: 4561084, BioRad), and transferred to 0.2 mm nitrocellulose membranes (catalog no: 1620112, BioRad). The blotting membranes were probed with anti MALT1 (#2494, 1:1000, Cell Signaling), Bcl10 (sc-5273, 1:500, Santa Cruz Biotechnology), CYLD (sc-74435, 1:500, Santa Cruz Biotechnology), GAPDH (sc-32233, 1:5000, Santa Cruz Biotechnology), CARD10/CARMA3 (sc-071849, 1:500, Santa Cruz Biotechnology), N4BP1 (ab133610, 1:1000, Abcam), CARD9 (sc- 374569, 1:500, Santa Cruz Biotechnology) antibodies. Bands were detected using the Western lightning ECL detection system (catalog no: 32106; Thermo Fisher Scientific).

### Macrophage chemotaxis

Macrophage migration assays were performed using the IncuCyte ClearView migration plate (Essen BioScience, catalog no.: 4582) coated with Matrigel (50 μg/mL). In brief, primary macrophages were isolated from WT, MALT1-PD or MALT1 KO mice as described above and were suspended in 30 μl of DMEM without FBS. Primary macrophages (3x 10^5^) were then placed in the top chamber of the IncuCyte migration plate, with or without MALT1 protease inhibitors (Mepazine or MLT-748, 5 μM) or a CARD9 inhibitor (BRD5529, 25 μM) in both the top and bottom well. GL261 (×10^5^) cells were placed in the bottom well of the IncuCyte migration plate, and migration of primary macrophages from the top to bottom well was measured every 2 hours over a period of 48 hours using the IncuCyte system and bottom-side phase analysis. Alternatively, 6x10^5^ RAW264.7 macrophages were seeded into an IncuCyte® Imagelock 96-Well Plate (Sartorius, catalog no.: BA-04855). After 24 h, a cell monolayer wound was created with IncuCyte® Woundmaker Tool (Sartorius, catalog no.: 4563). Next, the medium was replaced with GL261 conditioned media (GL261-CM) in the presence of indicated treatments. To prepare GL261-CM, GL261 tumor cells were seeded with 8 ml of serum free medium in a 25-cm dish, and the culture medium were collected after 4 hours followed by filtration using a 0.45-μm filter.

### Preclinical studies

#### GBM implantation in mice

Four to six weeks old C57BL/6 were used in this study. Mice were placed in a clear plexiglass anesthesia induction box that allowed unimpeded visual monitoring of the animals. Induction was achieved by administration of 2% isoflurane mixed with oxygen for a few minutes. Once the plane of anesthesia was established, it was maintained with 1-2% isoflurane in oxygen via a nose cone, and the mouse was transferred to the stereotaxic frame. GL261or CT2A cells were cultured to approximately 90% confluence and a total of 2 × 10^5^ cells in 2 μL DMEM without FBS was injected in the right striatum at a depth of 3.0 mm (coordinates regarding bregma: 2.5 mm lateral and 1.5 mm posterior) of C57BL/6 WT or MALT1-PD mice. Animals were euthanized 21 days following glioma cell implantation. All procedures used in the present study followed the Principles of Laboratory Animal Care from NIH (protocol number IS00020798).

For mouse drug treatment, on day 5 after surgery, tumor-bearing mice were randomized and treated with the MALT1 inhibitor mepazine (8mg/kg IP BID) for a total of 16 days as previously described^52^, alone or in combination with TMZ (25 mg/kg IP QD) for 5 days. Control animals were treated with the same volume and 10% DMSO.

### MRI

Imaging of mice was performed at the Rangos Research Center Animal Imaging Core at UPMC Children’s Hospital of Pittsburgh. All mice received general inhalation anesthesia with Isoflurane for *in vivo* brain imaging. *In vivo* MRI brain image was carried out using a Bruker BioSpec 70/30 USR spectrometer (Bruker BioSpin MRI) operating at 7-Tesla field strength, equipped with an actively shielded B-GA12S2 gradient system with 440 mT/m gradient strength and slew rate of 3440 T/m/s, as well as a quadrature radiofrequency volume coil with an inner diameter of 35 mm. T2-weighted anatomical and T1-weighted contrast multi-planar MRI of 11 to 21 slices (depending on tumor size and to cover the whole brain volume) were acquired with the following parameters: field of view (FOV) = 2.0 cm, matrix = 256 ′ 256, slice thickness = 0.6 mm, in-plane resolution = 78 µm ′ 78 µm. T2-weighted anatomical imaging was acquired with the RApid imaging with Refocused Echoes (RARE) with echo time (TE) = 12 msec, RARE factor = 8, effective echo time (TE) = 48 msec, repetition time (TR) = 1600 msec, and flip angle (FA) = 180°. Gadolinium (Gd) enhanced T1-weighted imaging was used to highlight the tumor volumes. Clinical grade Multihance (Gadobenate dimeglumine injection solution, 529 mg/ml, Bracco Diagnostics, Inc.) was administered subcutaneously with a 0.5 mmol/kg dosage. T1-weighted contrast imaging was acquired with the Fast Low Angle Shot (FLASH) sequence with 3 flip angles (FA): 10°, 20°, and 30°, TE = 2.349 msec, and TR = 92.590 msec. The images shown in the figures were acquired with FA = 30°. The multi-planar T1-weighted images were exported to DICOM format and analyzed by blinded independent observers using the open-source ITK-SNAP (http://www.itksnap.org) brain segmentation software.

### Flow cytometry for TME characterization

The entire tumor was removed, and samples were prepared for flow cytometry. Tumor tissues were homogenized for 15 minutes at 37°C in 3.2 mg/mL collagenase type IV, 2 mg/mL soybean trypsin inhibitor, and 1.0 mg/mL deoxyribonuclease I (Worthington Biochemical). After digestion, cell suspensions were passed through a 70-μm filter, and collagenase was inactivated with 0.05 M EDTA in PBS (pH 7.4). Cells were harvested twice by centrifugation in PBS at 400×g for 6 min. About 1 × 10^6^ cells were blocked with Rat Anti-Mouse CD16/CD32 (Mouse BD Fc Block, catalog no.: 553141) for 15 min before staining with specific labeled antibodies (Supplementary Table 1). The data were acquired using a Cytek Aurora and analyzed using Flow Jo software. The gating strategy is shown in Figure S10.

### Single Cell Filtering, Normalization, Clustering

GL261 tumors were enzymatically and mechanically dissociated into a single cell suspension as described above. Live cells were further isolated from the single-cell suspension using the Dead Cell Removal Kit (Miltenyi Biotec). Cell viability was confirmed prior to the start of cell-hashing with acridine-orange-propidium iodide stain (Nexcelcom Biosciences) and a Cellometer Auto 2000 Viability Counter (Nexcelcom Biosciences). Three tumors were cell-hashed together into a single suspension using TotalSeq C Antibodies (Biolegend). Library Preparation (Chromium Next GEM Single Cell 3’ Reagent kit v3.1) was conducted as per the manufacture’s protocol. The quality of the cDNA libraries was verified using a High Sensitivity D5000 ScreenTape and 4200 Tapestation system (Agilent Technologies). Samples were sequenced on an Illumina Novaseq 6000 PE150 (Read 1 150bp, i7 Index 10bp, i5 Index 10bp, Read 2 150bp). Cell Ranger (10x Genomics) analysis pipeline was used to perform alignment, filtering, and counting of barcodes and unique molecular identifiers. The sequencing data was aligned to the mouse genome mm10. Aggregated data matrices for WT & MALT1-PD were imported using the Seurat package^57^ in R and filtered by features including at least 3 cells and including cells which presented at least 200 features, then the datasets were merged and normalized together via *NormalizeData* in the Seurat pipeline, resulting in 42712 remaining cells in the dataset, of which 21411 were WT and 21301 were MALT1-PD. Data was then scaled using the Seurat *ScaleData* function and the first 30 components of the principal component analysis were used to construct the UMAP. Seurat *FindClusters* was used with a resolution of 0.5 to determine the 26 clusters within the dataset. Clusters were annotated using a combination of top differential genes per cluster and unbiased cell type recognition using the SingleR package with the mouseRNAseq dataset from the celldex package. Differentially expressed genes were determined between WT and PD groups using Suerat *FindMarkers* with the Mast algorithm as the test. Only genes expressed in at least 25% of the cells being compared were used, and a cutoff for the top genes was set to a foldchange ≥ 2 and a p-value ≤ 0.05, with a cutoff of the top 50 most significant genes by p value if a list were to contain more than 50 significant genes. Heatmaps show the log2 average expression per either WT or PD cells in the data per genes. Top differential genes per groups of cells were utilized as input to the Ingenuity Pathway Analysis software, with foldchange being the primary input variable and utilizing only datasets with mouse as the primary species.

### Open field analysis

Locomotor activity was evaluated using a video tracking system. 20 days following tumor implantation, animals were placed inside an Open Field apparatus. Open field exploration was carried out on a 44 × 44 cm arena, surrounded by transparent walls 30 cm high. The floor of the arena was divided into nine equal squares by black lines. Mice were placed in the left rear quadrant of the apparatus. Each of the movement types, i.e., resting and slow-moving, was recorded for 5 min following 15-second adaptation period, and the ratio was analyzed using the Smart V3.0 software.

### Bioinformatic analyses of public databases

The Cancer Genome Atlas (TCGA) was explored via the Gliovis platform (http://gliovis.bioinfo.cnio.es/)^12^. Differential expression of *MALT1* in human GBMs was performed using single cell RNA-sequencing (scRNAseq) data obtained by Darmanis *et al.,* made publicly available on the Gene Expression Omnibus (GEO) GSE84465 dataset^33^. This dataset includes 3589 cells from 4 IDH wild type glioblastomas. The authors used antibody-mediated cell sorting, RNA cluster-based sorting, and copy-number variation analysis to identify seven cell types: immune, oligodendrocytes, oligodendrocyte precursors (OPCs), vascular, neurons, astrocytes, and neoplastic cells. Full details of tumor collection, RNA-sequencing, and quality control parameters can be found in the original paper^33^. Gene counts, cell type phenotyping and 2D-tSNE (t-distributed stochastic neighbor embedding) representation of included cells were downloaded from http://gbmseq.org/. MALT1 plots in non-tumor versus GBM in different cells subsets were prepared using bulk RNA-seq data from Klemm et al.^35^ (https://joycelab.shinyapps.io/braintime/). Comparison of MALT1 mRNA expression in 3 different GBM molecular subtypes was evaluated as previously described^58^. Probability of survival of 150 patients diagnosed with GBM were grouped by high (top 50%) or low (bottom 50%) MALT1 mRNA level^59^.

### Statistics and reproducibility

All statistical analyses were performed using GraphPad Prism 9 software. Statistical significance was calculated by two-tailed paired or unpaired Student’s t-tests on two experimental conditions and one-way or two-way analysis of variance (ANOVA) test when more than two experimental groups were compared. P values were adjusted for multiple comparisons by Tukey method and a P value of <0.05 was defined as statistically significant. For *in vitro* experiments, no data was excluded, and the experiments were performed with at least with two to three biological replicates. Data distribution was assumed to be normal. For *in vivo* studies, tumor measurement and analysis were performed blindly by different researchers to ensure that the studies were run in a blind manner with at least five animals allocated per group. Animals were excluded only if they died or had to be killed according to protocols approved by the animal experimental committees before the end point. Sample sizes in this study were estimated based on previous experience that showed significance. All animals were randomized and exposed to the same environment. Survival data were collected blinding and estimated by a log-rank (Mantel–Cox) test was used. All the results are shown as mean ± SD. The exact sample sizes are indicated in the figure legends. To analyze the relationship between MALT1 expression and survival of patients with GBM, data were downloaded from GlioVis (http://gliovis.bioinfo.cnio.es/). Patients were divided into low and high groups, and long–rank survival analysis was performed with GraphPad Prism 9 to generate Kaplan–Meier survival curves.

## Acknowledgments

The authors would like to thank Dr. Baoli Hu from University of Pittsburgh for providing the glioblastoma cell lines. The authors thank all members of the Lucas/McAllister labs for advice and support. This project involved use of the Animal Imaging, Flow cytometry and Histology core facilities of the University of Pittsburgh Department of Pediatrics, Rangos Research Center. This work utilized the UPMC Hillman Cancer Center Biostatistics Facility, a shared resource at the University of Pittsburgh supported by the CCSG P30 CA047904.

## Funding

NIH 1K99NS135130-01 (JHA)

UPMC Hillman Innovative Postdoctoral Scholar Award (JHA)

UPMC Children’s Hospital of Pittsburgh Research Advisory Committee (RAC) award (JHA) University of Pittsburgh/Carnegie Mellon University Medical Scientist Training Program (HB) Conover Scholar award (HB)

NIH F30 CA284607-01 (HB) NIH 5K12HD052892 (LMM)

UPMC Hillman Cancer Center Support Grant 5P30CA047904 (LMML).

## Author contributions

Conceptualization: JHA, PCL, LMML

Methodology: JHA, GND, SSY, HC, LMM, CS, KES, RA, YLW, GK, RB, MS, PE, DH, AM

Investigation: JHA, SSY, GND, HC LMM Visualization: JHA, SSY, GND, HC, LMM Breeding: LK

MALT1-PD/KO mouse generation: PJG, JB Funding acquisition: JHA, PCL, LMML Project administration: PCL, LMML Supervision: PCL, LMML

Writing – original draft: JHA

Writing – review & editing: JHA, PCL, LMML.

## Data and materials availability

All data associated with this study are present in the paper or the Supplementary Materials. Single cell RNA-seq data will be deposited into the National Center for Biotechnology Information (NCBI) Gene Expression Omnibus database. All noncommercially available new materials will be made available to nonprofit or academic requesters upon completion of a standard material transfer agreement. Requests for materials may be made by contacting LMML.

