## Supplementary Material for "MALT1 protease inhibition restrains glioblastoma progression by reversing tumor-associated macrophage-dependent immunosuppression"

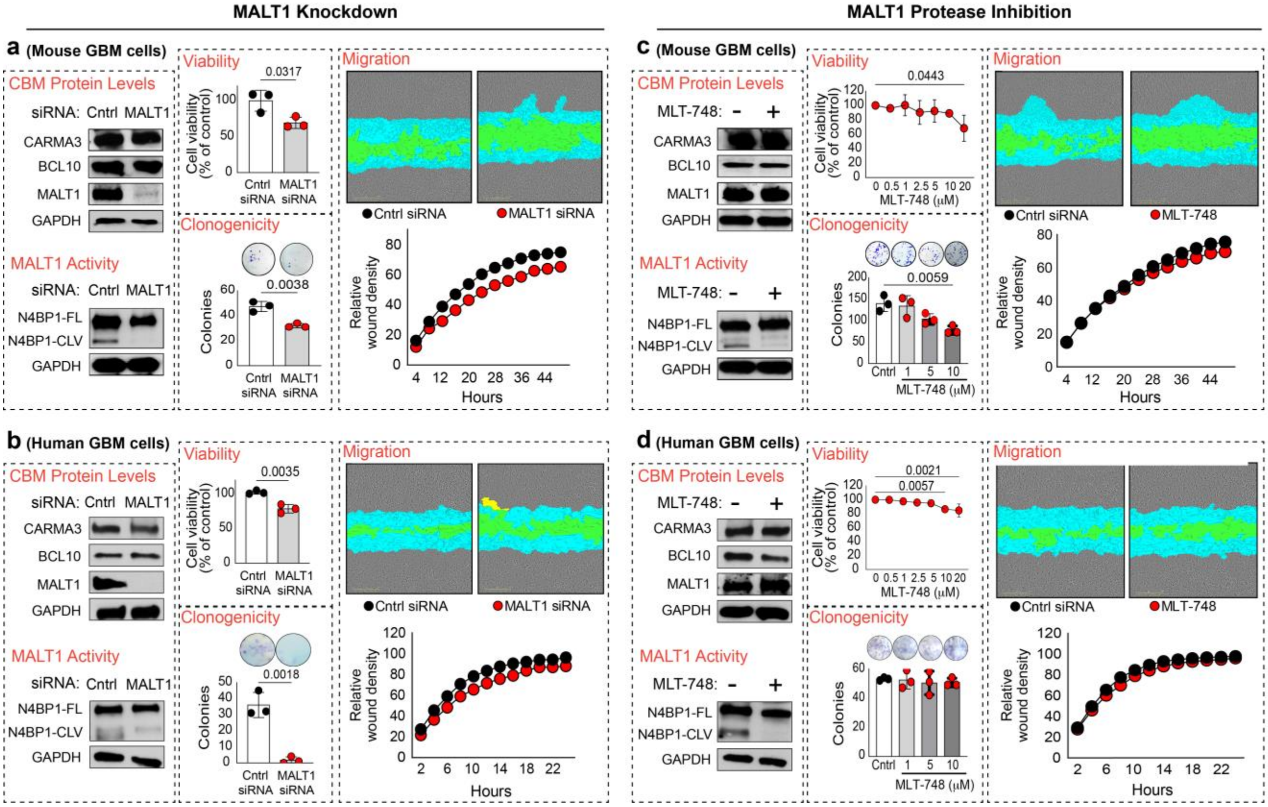


**Supplementary Figure 1. Effects of MALT1 blockade on GBM tumor cell behavior. (a)** CBM components are present in mouse GBM cells (GL261 line), as determined by Western blotting. MALT1 shows constitutive protease activity, cleaving the substrate N4BP1, and siRNA-mediated knockdown of MALT1 abrogates N4BP1 cleavage as assessed by a reduction in the cleaved N4BP1 product on Western blot. MALT1 knockdown for 72 hours significantly reduces GL261 cell viability and clonogenic potential but has no significant effect on cell migration in the scratch wound assay. **(b)** Similar to what we observed with the mouse GL261 line (panel a), CBM components are present and active in human GBM cells (U87MG line), and MALT1 knockdown abrogates the constitutive MALT1 protease activity. MALT1 knockdown for 72 hours significantly reduces U87MG cell viability, dramatically impairs clonogenic potential, but has no significant effect on cell migration in the scratch wound assay. **(c)** MALT1 protease inhibition with MLT-748 (5 µM for two days) effectively blocks NFBP1 cleavage in mouse GL261 cells and does not alter the expression level of any CBM complex components, as assessed by Western blotting. MLT-748 reduces GL261 cell viability and clonogenic potential but does not affect cell migration. **(d)** MALT1 protease inhibition with MLT-748 (5 µM for two days) effectively blocks NFBP1 cleavage in human U87MG cells and does not alter the expression level of any CBM complex components, as assessed by Western blotting. MLT-748 has a small negative effect on U87MG cell viability, but no impact on clonogenicity or cell migration. All experiments were repeated at least three times. Quantification of scratch wound closure is plotted as a continuous function of time, with representative end-point images shown above each plot. All values are represented as mean ± SD. Data were analyzed by 1-way ANOVA, followed by Tukey's multiple-comparisons test or unpaired, 2-tailed Student's t test. Significant P values are indicated in the figure.

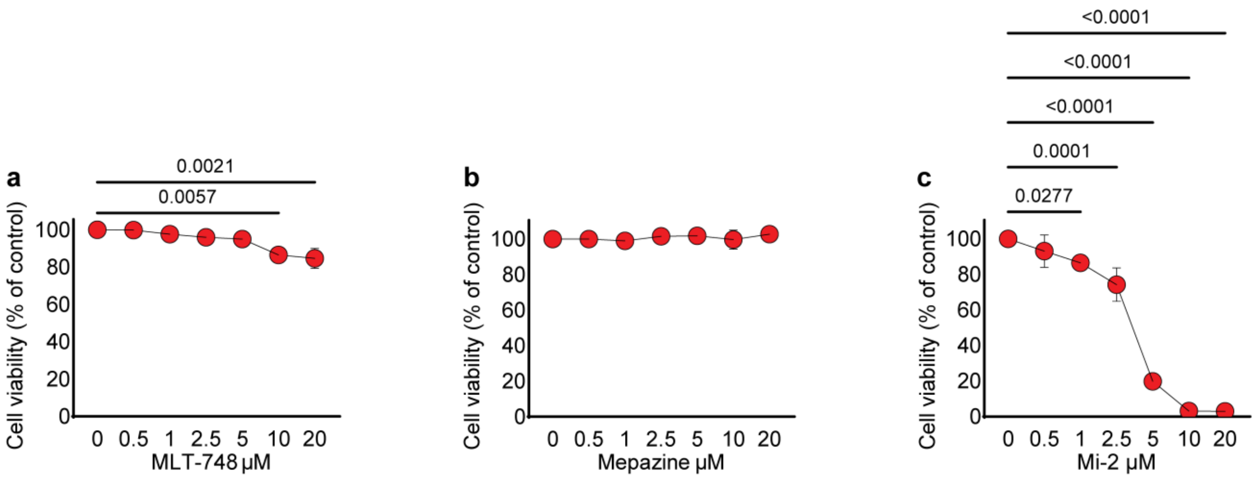


**Supplementary Figure 2. Compound MI-2 inhibits the growth of human GBM cells.** Effects of three different MALT1 protease inhibitors, **(a)** MLT-748, **(b)** Mepazine, and **(c)** MI-2, on human U87MG GBM cell viability. Cells were treated with compounds at the indicated concentrations for 3 days. Results represent an average of 3 independent experiments. Graphs show the percentage of viable cells relative to DMSO vehicle control. Data represent the mean ± SD. Data were analyzed by 1-way ANOVA, followed by Tukey's multiple-comparisons test. Significant p values are indicated in the figure.

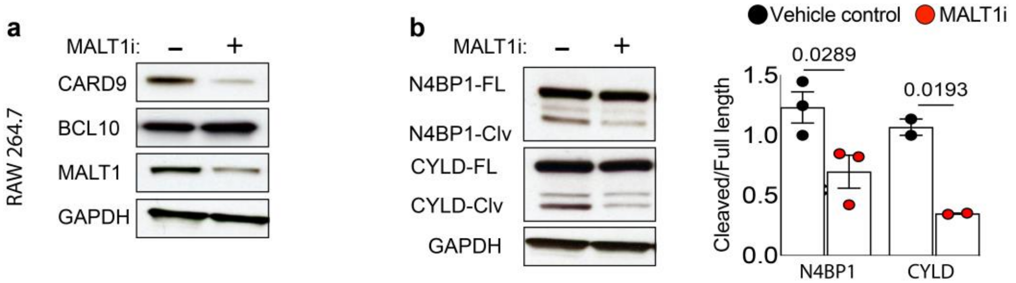


**Supplementary Figure 3. CBM components (CARD9-BCL10-MALT1) are present and MLT-748 inhibits MALT1-protease activity in macrophages**. **(a)** Mouse Macrophage (RAW264.7) cells were treated with or without 5 μM MLT-748 (MALT1i) for 2 days before harvesting and immunoblot analysis. Experiments were performed in triplicate. (**b**) The reduction in cleavage of MALT1 proteolytic substrates, N4BP1 and CYLD, confirms MALT1 protease inhibition. Western blot results are quantified by densitometry as shown in the bar graphs to the right. Data are shown as mean ± SD. Statistical significance was determined by two-tailed Student's t-test; P values <0.05 are shown in the figure.

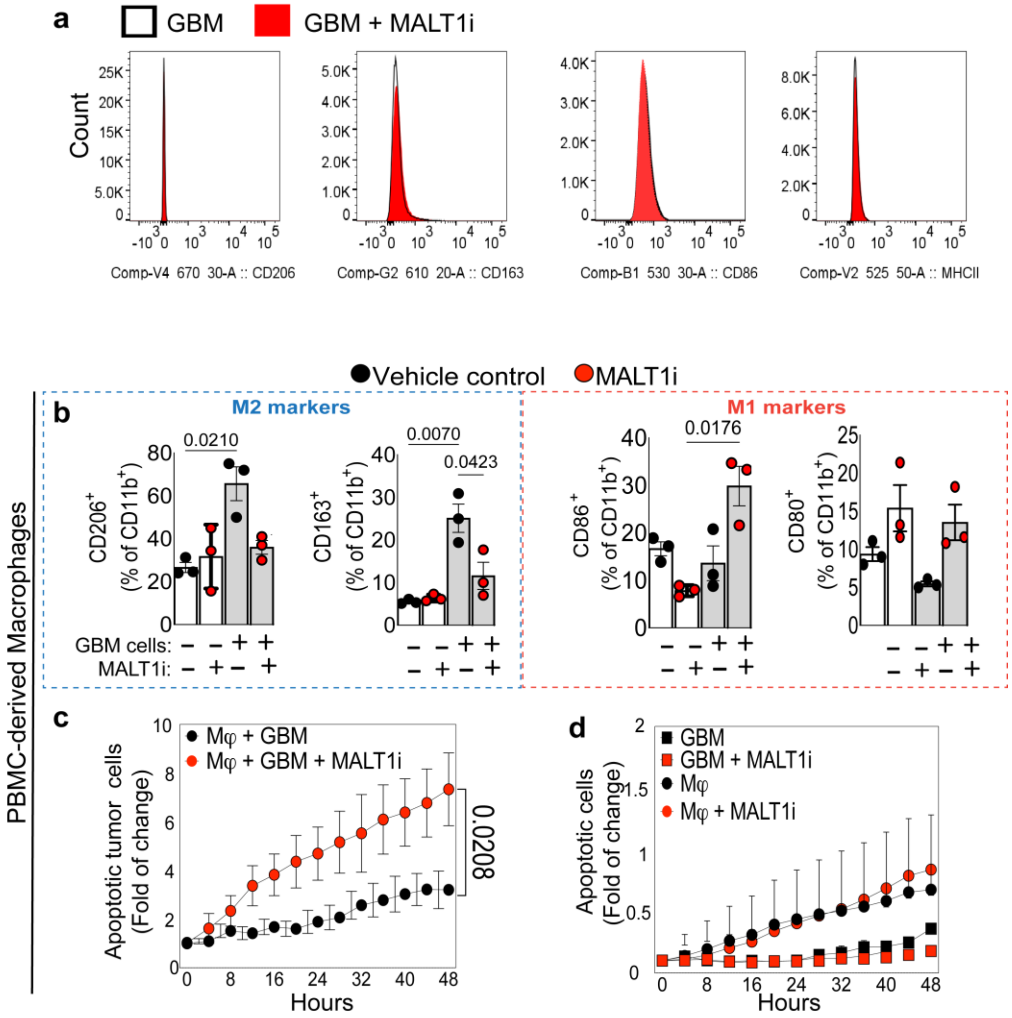


**Supplementary Figure 4. Additional control experiments for confirming that inhibition of MALT1 protease reprograms M2-like GBM-associated macrophages into M1-like macrophages with enhanced anti-tumor activity. (a)** Pharmacologic MALT1 protease inhibition has no effect on expression of CD206, CD163, CD86 and MHCII in GL261 GBM cells. Cells were treated -/+ 5 µM MLT-748 for 2 days. Phenotypic markers were analyzed by flow cytometry gating on CD11b negative cells. The Figure shows representative histograms. **(b)** Changes in M2/M1-like polarization markers after Human primary PBMC-derived macrophages were cocultured with U87MG human GBM cells in the absence or presence of MALT1 inhibitor (MALT1i; MLT-748, 5 µM) for 48 hours. **(c.d)** The Incucyte® live-cell analysis system was used to measure tumor cell killing in real-time. A dual color monitoring system tracked tumor cell killing by using fluorescently labeled tumor cells (red-Cell Tracker) and caspase-3/7 reagent (green) to track apoptosis. **(c)** Co-cultures of human primary macrophages with human U87MG-GBM cells were performed to compare tumor cell apoptosis in the absence/presence of MALT1i. **(d)** Controls demonstrating that MALT1i has no effect on apoptosis of U87MG-GBM tumor cells or on primary human macrophage when these cells are cultured alone. All values are represented as mean ± SD. Data were analyzed by 1-way ANOVA, followed by Tukey's multiple-comparisons. Results are representative of 3 independent experiments in duplicate, using 3 different donors. Significant p values are indicated in the figure.

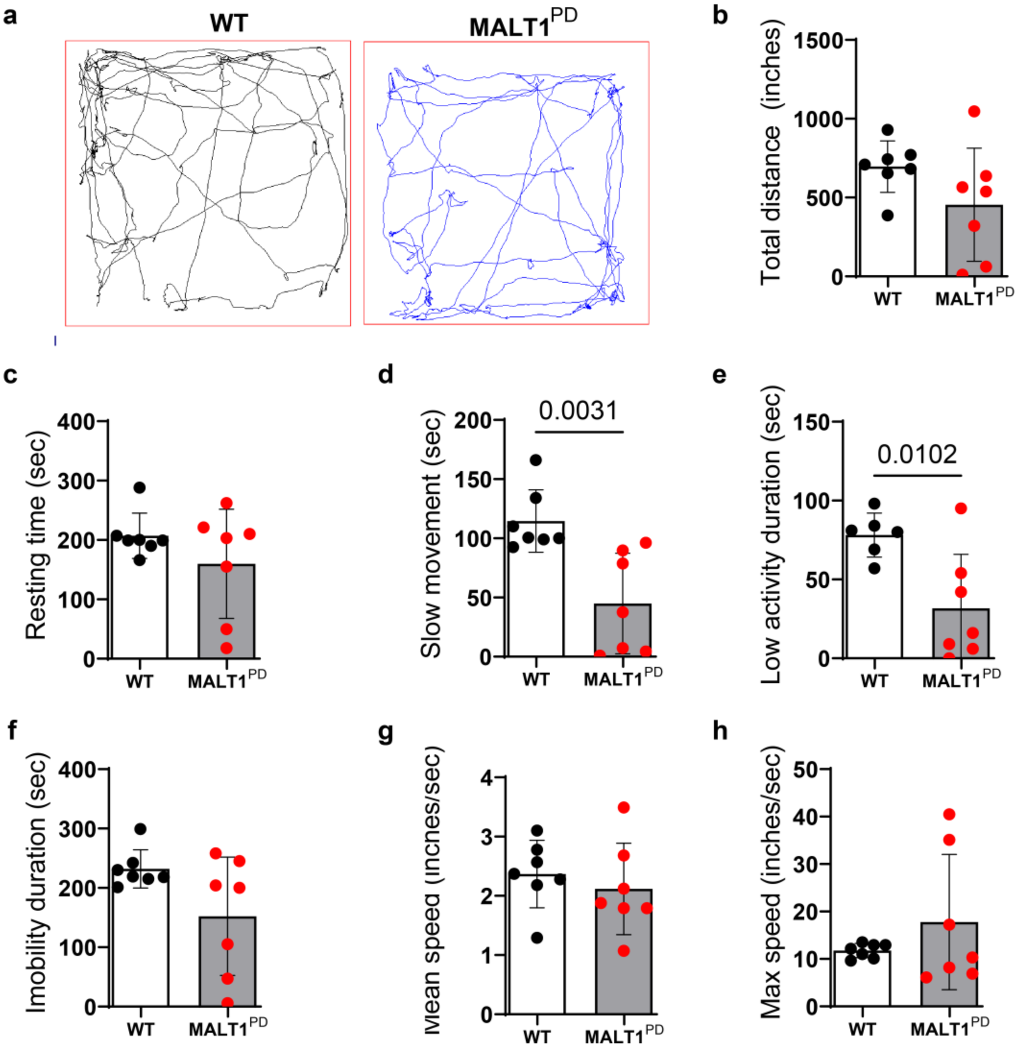


**Supplementary Figure 5. MALT1-PD mice bearing GBM GL261 tumors demonstrate improved locomotor ability in comparison to WT controls.** The following parameters were analyzed: **(a)** Representative tracks of control WT (n=7) and MALT1-PD mice (n=7) in the open field chamber over 5 min, **(b)** total distance traveled, **(c)** resting time, **(d)** time spent in slow movement, **(e)** low activity time, **(f)** immobility, **(g)** mean speed and **(h)** maximum speed. Statistical analysis was performed using an unpaired t-test. Data are displayed as mean ± SD. Significant P values are indicated in the figure.

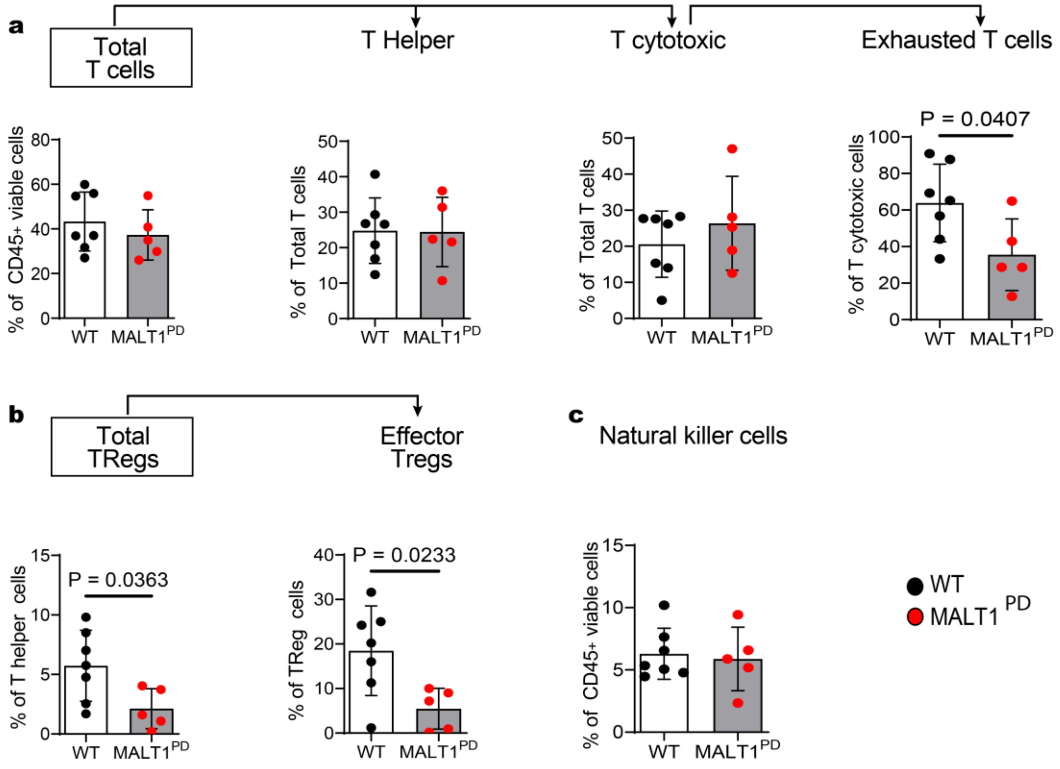


**Supplementary Figure 6. Lack of MALT1-protease activity in the host TME results in less lymphocyte exhaustion and less Treg in the GBM TME. (a)** Quantification of the percentages of total T lymphocytes ($CD3^+$), T helper ($CD3^+CD4^+CD8^-$), T cytotoxic ($CD8^+CD3^+CD4^-$) and Exhausted T cytotoxic ($CD3^+CD8^+CTLA\text{-}4^+TIM3^+LAG\text{-}3^+PD\text{-}1^+$), **(b)** Treg ($CD4^+CD25^+FOXP3^+$), eTreg ($CD4^+CD25^+FOXP3^+PD1^+CTLA\text{-}4^+$) and **(c)** Natural killer ($NK1.1^+$) cells. Data are shown as mean ± SD from two independent experiments. Statistical significance was determined by two-tailed Student's t-test; significant P values are indicated in the figure. Each dot represents an independent mouse.

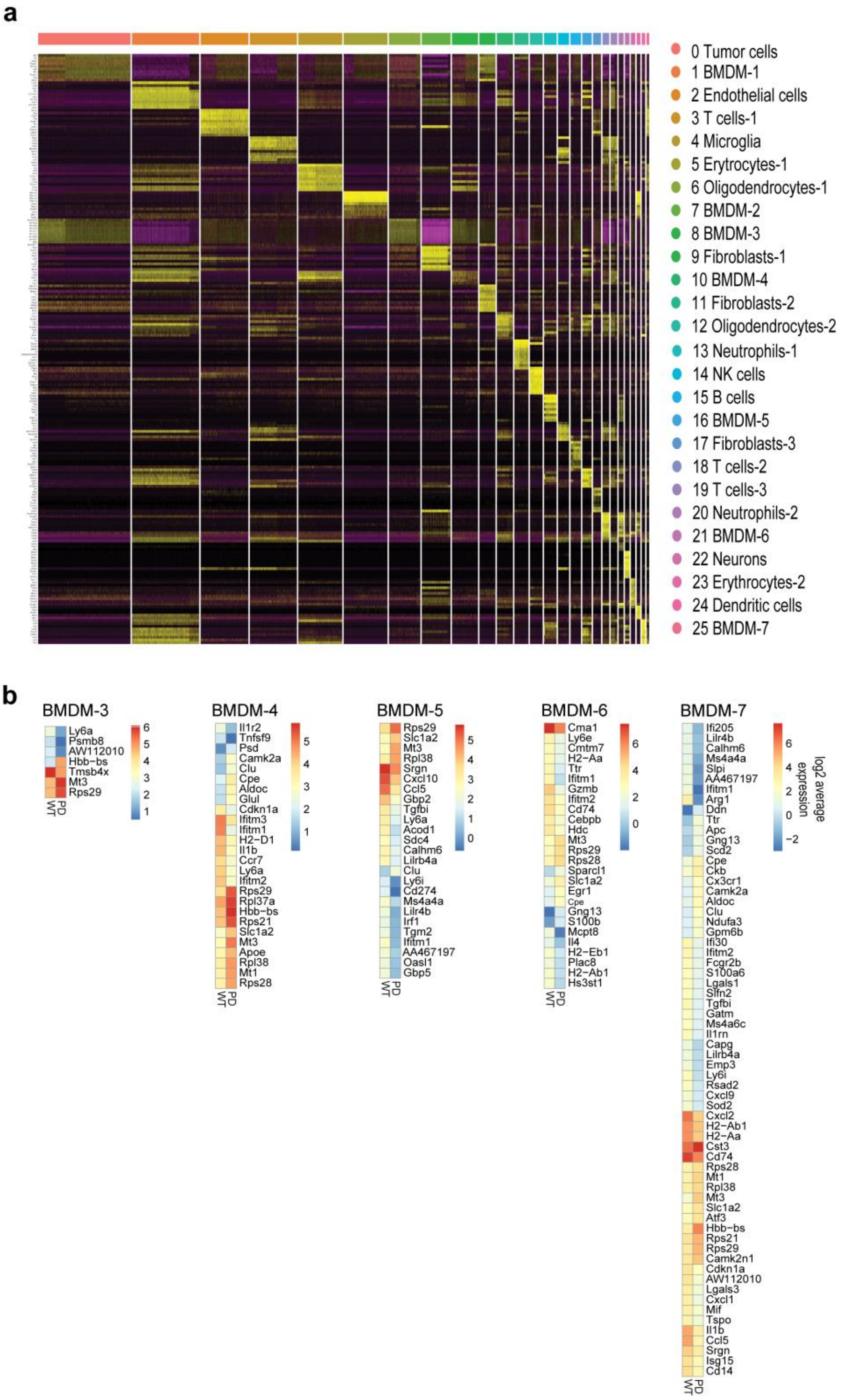


**Supplementary Figure 7. Single cell RNA-Seq clustering and gene expression patterns. (a)** Heatmap representation of the ten most highly expressed genes within each of the 26 clusters present in the GBM tumors in mice. Hierarchical clustering of cells grouped by top expression of genes. The colors indicated on the legend to the right also correspond to the color legend provided in Figure 4. **(b)** Heatmap representation of differentially expressed genes in bone marrow derived macrophage (BMDM) clusters 3-7 from GBM-bearing WT vs MALT1-PD mice. Heatmaps reflect log2 average expression of top differentially expressed genes in WT versus MALT1-PD cells within each macrophage cluster. Average-scaled expression is indicated on the color gradient.

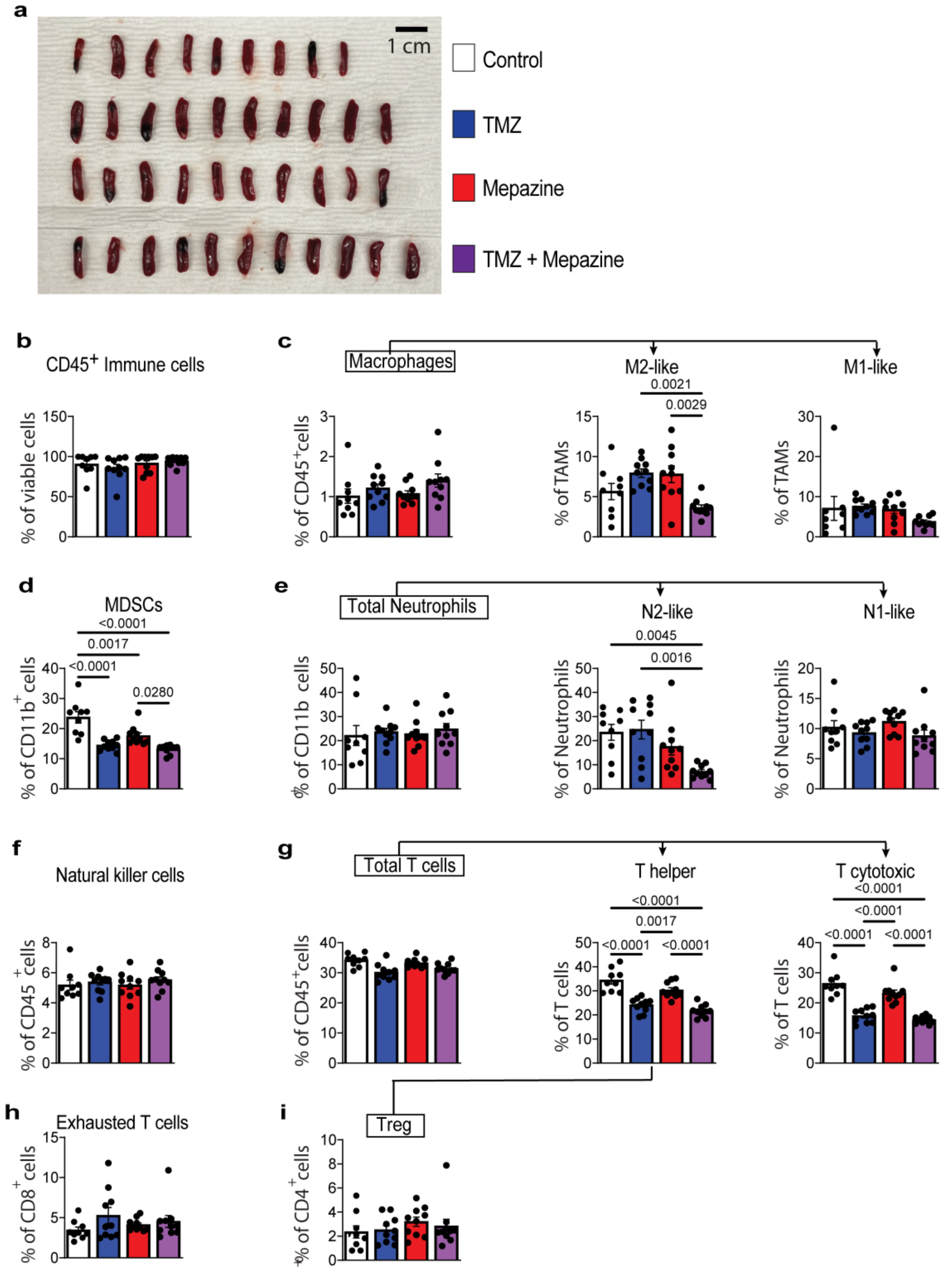


**Supplementary Figure 8. Flow cytometry analyses of spleens from GBM-bearing mice.** **(a)** Photo of the spleens of mice undergoing treatment as described in Fig. 7a. **(b-i)** flow cytometry analysis of spleen cells showing the percentage of **(b)** Total immune cells ($CD45^+$), **(c)** macrophages ($CD11b^+F480^+$), M2-like macrophages ($CD11b^+F480^+CD206^+Arg1^+$) and M1-like-macrophages ($CD11b^+F480^+iNos^+MHCII^+$), **(d)** MDSCs ($CD11b^+Ly6C^+Ly6G^-$), **(e)** Neutrophils ($CD11b^+Ly6G^+Ly6C^-$), N2-like Neutrophils ($Arg1^+CD206^+$), N1-like Neutrophils ($iNos^+Arg1^-$), **(f)** Natural killer cells ($NK1.1^+$), **(g)** T lymphocytes ($CD3^+$), T helper ($CD3^+CD4^+CD8^-$), T cytotoxic ($CD8^+CD3^+CD4^-$)**, (h)** Exhausted T cells ($CD3^+CD8^+CTLA\text{-}4^+TIM3^+LAG\text{-}3^+PD\text{-}1^+$) and **(i)** Treg ($CD4^+CD25^+FOXP3$). Data are shown as mean ± SD. n = 9 for control, n=9 for TMZ, n=9 for MALT1i and n=10 TMZ + MALT1i. Statistical significance was determined by one way ANOVA followed by Tukey; significant P values are indicated in the figure.

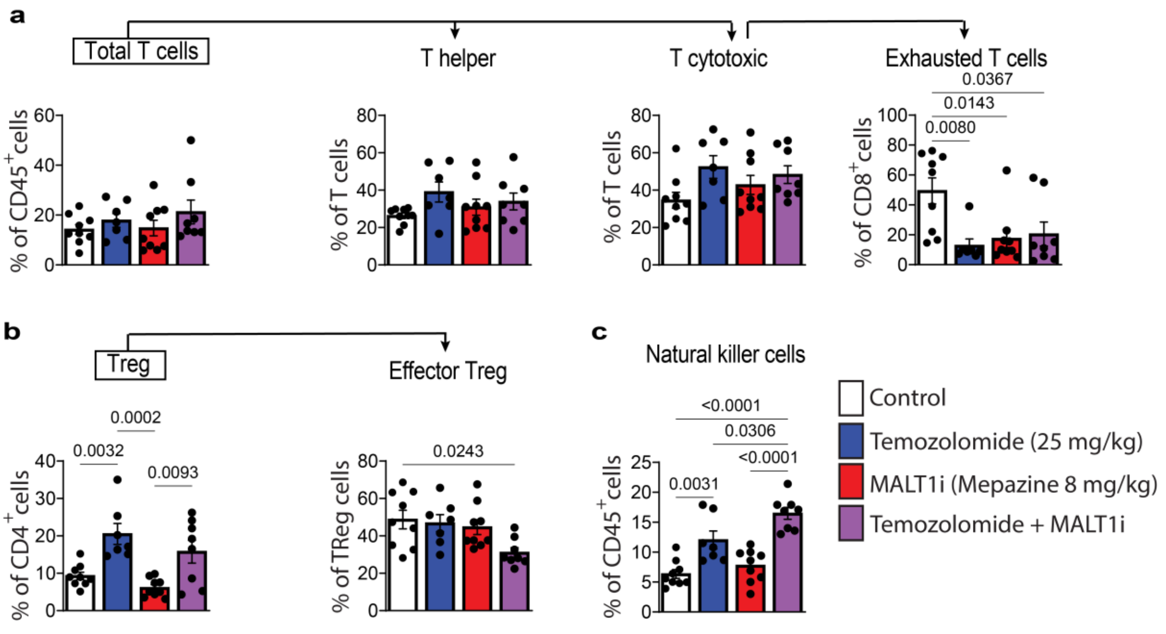


**Supplementary Figure 9. Effects of mepazine and TMZ on lymphocytes in the GBM TME.** Mouse GL261 cells were transplanted into mouse brains through intracranial injection to establish GBM tumors. Five days after transplantation, tumor-bearing mice were treated with vehicle control (DMSO 5%), TMZ (25 mg/kg for 5 days), Mepazine (MALT1i) (8 mg per kg/twice per day for 16 days) or TMZ. Mouse brains bearing GBM tumors were collected for further analyses on day 21. **(a)** Quantification of the percentages of total T lymphocytes ($CD3^+$), T helper ($CD3^+CD4^+CD8^-$), T cytotoxic ($CD8^+CD3^+CD4^-$), Exhausted T cytotoxic($CD3^+CD8^+CTLA\text{-}4^+TIM3^+LAG\text{-}3^+PD\text{-}1^+$), **(b)** Treg ($CD4^+CD25^+FOXP3^+$), eTreg ($CD4^+CD25^+FOXP3^+PD1^+CTLA\text{-}4^+$) and **(c)** Natural killer($NK1.1^+$). Data are shown as mean ± SD. Statistical significance was determined by One-way ANOVA followed by post-hoc Tukey; significant P values are indicated in the figure. Each dot represents an independent mouse.

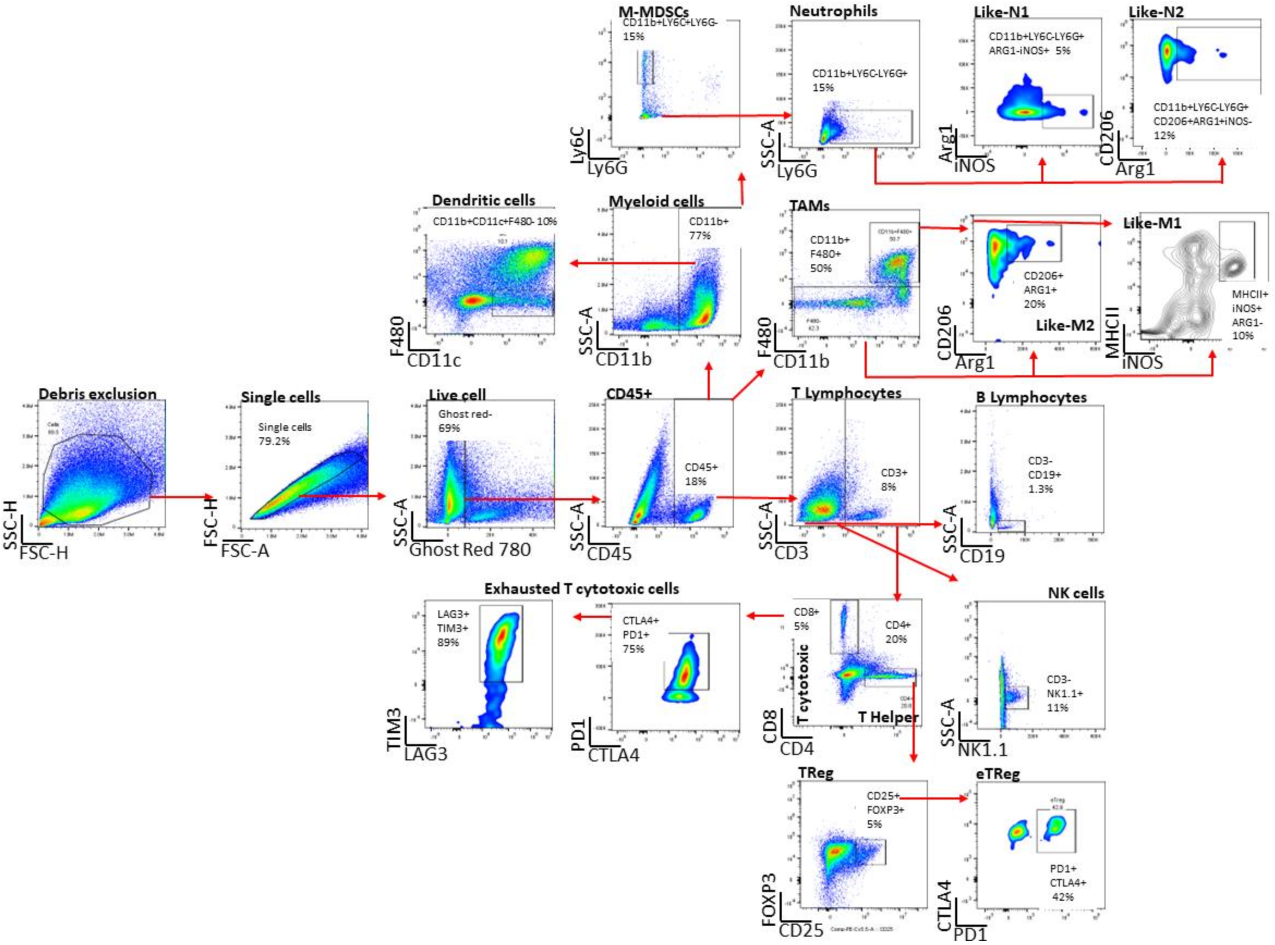


**Supplementary Figure 10.** Gating strategy.

**Table S1.** List of antibodies and antibody dilutions used in the described experiments.

| | Marker | Clone | Target cells | Fluorochrome | Company | Dilution |
|---|---|---|---|---|---|---|
| | ***In vitro* - Human** | | | | | |
| 1 | CD206 | 15-2 | M2-like | PE/Dazzle594 | eBioscience, cat. No. 321130 | 1:20 |
| 2 | CD14 | 63D3 | Myeloid cells | BV750 | eBioscience, cat. No. 367136 | 1:20 |
| 3 | MHCII | LN2 | M1-like | APC | eBioscience, cat. No. 32812 | 1:20 |
| 4 | CD163 | GH1/61 | M2-like | BV421 | eBioscience, cat. No. 333612 | 1:20 |
| 5 | CD86 | IT2.2 | M1-like | FITC | eBioscience, cat. No. 305414 | 1:20 |
| 6 | CD80 | 2D10 | M1-like | BV605 | eBioscience, cat. No. 305225 | 1:20 |
| | ***In vitro*- Mice** | | | | | |
| 1 | *CD206* | C068C2 | M2-like | BV650 | BioLegend, cat. No. 141723 | 1:33 |
| 2 | *CD11b* | M1/70 | Myeloid cells | BV750 | eBioscience, cat. No. 101267 | 1:500 |
| 3 | *MHCII* | M5/114 | M1-like | BV510 | eBioscience, cat. No. 107636 | 1:100 |
| 4 | *CD163* | S150491 | M2-like | PE-Dazzle594 | eBioscience, cat. No. 155316 | 1:100 |
| 5 | *CD86* | GL-1 | M1-like | FITC | eBioscience, cat. No. 105018 | 1:100 |
| 6 | *CD80* | 16-10ª1 | M1-like | BV605 | eBioscience, cat. No. 104729 | 1:300 |
| | ***In vivo*** | | | | | |
| 1 | CD45 | 30-F11 | Pan marker | BUV395 | BD Bioscience, cat. No. 565967 | 1:100 |
| 2 | CD3 | 17A2 | T cells | BUV496 | BD Bioscience, cat. No. 741117 | 1:100 |
| 3 | CD4 | RM4-5 | T helper cells | BUV563 | BD Bioscience, cat. No. 741217 | 1:80 |
| 4 | CD8 | 53-67 | Cytotoxic T cells | BUV615 | BD Bioscience, cat. No. 613004 | 1:200 |
| 5 | CD25 | PC61.5 | Tregs | PE-Cy5.5 | eBioscience, cat. No. 35-0251-82 | 1:80 |
| 6 | Foxp3 | FJK-16s | Tregs | PE | eBioscience, cat. No. 12-5773-82 | 1:20 |
| 7 | NK1.1 | PK136 | NK cells | PE-Cy7 | eBioscience, cat. No. 25-5941-82 | 1:80 |
| 8 | CD19 | eBio1D3 (1D3) | B cells | BUV661 | BD Bioscience, cat. No. 612971 | 1:80 |
| 9 | Ly6C | HK1.4 | monocytes | PE/Dazzle594 | BioLegend, cat. No. 128044 | 1:333 |
| 10 | Ly6G | 1A8 | neutrophils | BUV805 | BD Bioscience, cat. No. 741994 | 1:100 |
| 11 | F4/80 | BM8 | macrophages | PerCP-Cy5.5 | eBioscience, cat. No. 45-4801-82 | 1:40 |
| 12 | CD11b | M1/70 | myeloid cells | BV570 | BioLegend, cat. No. 101233 | 1:20 |
| 13 | MHCII | M5.114/15.2 | dendritic cells | BV510 | BioLegend, cat. No. 107636 | 1:100 |
| 14 | CD11c | N418 | dendritic cells | SBlue 550 | BioLegend, cat. No. 117366 | 1:200 |
| 15 | PD-1 | 29F.1A12 | T cells | BV421 | BioLegend, cat. No. 135221 | 1:160 |
| 16 | CTLA4 | UC10-4B9 | T cells | BV605 | BioLegend, cat. No. 106323 | 1:20 |
| 17 | TIM3 | RMT3-23 | T cells | BV711 | BioLegend, cat. No. 119727 | 1:80 |
| 18 | LAG-3 | C9B7W | T cells | BV785 | BioLegend, cat. No. 125219 | 1:27 |
| 19 | iNOS | CXNFT | M1-like | FITC | eBioscience, cat. No. 53-5920-82 | 1:200 |
| 20 | CD206 | C068C2 | M2-like | BV650 | BioLegend, cat. No. 141723 | 1:33 |
| 21 | Arginase | A1exF5 | MDSC | AFluor 700 | eBioscience, cat. No. 56-3697-82 | 1:160 |
| 22 | PD-L1 | MIH5 | T cells | APC | BD Bioscience, cat. No. 564715 | 1:100 |
